# Redefining the Limits of Functional Continuity in the Early Evolution of P-Loop NTPases

**DOI:** 10.1101/2024.09.17.613540

**Authors:** Andrey O. Demkiv, Saacnicteh Toledo-Patiño, Encarni Medina Carmona, Andrej Berg, Gaspar P. Pinto, Antonietta Parracino, Jose M. Sanchez-Ruiz, Alvan C. Hengge, Paola Laurino, Liam M. Longo, Shina Caroline Lynn Kamerlin

**Affiliations:** Department of Chemistry – BMC, Uppsala University, BMC Box 576, S-751 23 Uppsala, Sweden; Protein Engineering and Evolution Unit, Okinawa Institute of Science and Technology, Graduate University (OIST), Okinawa, 904-0495, Japan; Departamento de Química Física, Facultad de Ciencias, Unidad de Excelencia de Química aplicada a Biomedicina y Medioambiente (UEQ), Universidad de Granada, 18071 Granada, Spain; Department of Chemistry and Biochemistry, Utah State University, Logan, Utah 84322-0300, United States; Institute for Protein Research, Osaka University, Suita, Japan; Blue Marble Space Institute of Science, Seattle, Washington 98154, United States; Earth-Life Science Institute, Tokyo Institute of Technology, Tokyo, 152-8550, Japan; School of Chemistry and Biochemistry, Georgia Institute of Technology, 901 Atlantic Drive NW, Atlanta, GA 30332, USA

**Keywords:** Walker A motif, P-Loop NTPase, phosphate binding loop, molecular fossil, primitive proteins

## Abstract

At the heart of many nucleoside triphosphatases is a conserved sequence motif that binds phosphate. A current model of early enzyme evolution proposes that this 6-8 residue motif could have sparked the emergence of the very first nucleoside triphosphatases – a striking example of evolutionary continuity from simple beginnings, if true. To test whether this provocative evolutionary model holds for the ancient and ubiquitous P-Loop NTPases, the properties of seven disembodied Walker A-derived peptides were extensively characterized by Hamiltonian replica exchange molecular dynamics simulations. Although dynamic flickers of nest-like conformations were observed, significant structural similarity between the situated peptide and its disembodied counterpart was not detected – even in the presence of orthophosphate or a nucleotide. Simulations suggest that phosphate binding is non-specific, with a slight preference for GTP over orthophosphate. Control peptides with the same amino acid composition but different sequences and situated conformations behaved similarly to the Walker A peptides with respect to conformational dynamics and phosphate binding, revealing no indication that the Walker A sequence is privileged as a disembodied peptide. We conclude that the evolutionary history of the P-Loop NTPase family is unlikely to have started with a disembodied Walker A peptide in an aqueous environment. The limits of evolutionary continuity for this protein family, and the environmental context within which it emerged, must be reconsidered. Finally, we argue that motifs such as the Walker A motif may represent incomplete or fragmentary molecular fossils – the true nature of which have been eroded by time.

**Significance Statement:** The first proteins were undoubtedly small, but when did those seeds emerge, and what did they look like? It is widely believed that the Walker A P-loop motif is a seed for the emergence of phosphate binding proteins, snugly binding phosphate in a structurally conserved nest. We probe this hypothesis through detailed computational characterization of disembodied Walker A and control peptides, showing that any nest formation is transient, and phosphate binding is weak and non-specific. Thus, we do not find structural continuity represented in the conserved Walker A motif, and current models of early P-loop evolution require revision. Further, care is required when interpreting highly conserved sequence fragments more broadly, as these may merely represent eroded molecular fossils.

## Introduction

P-Loop NTPases are the archetypical ancient protein family: Ubiquitously distributed across the tree of life, these domains have been recruited to catalyze phosphorylation reactions and to couple large molecular motions to the synthesis or hydrolysis of condensed phosphates (1–5). At the heart of the P-Loop fold lies a primitive active site in which the *α*- and *β*-phosphate moieties of an NTP molecule rest atop a crown or nest of backbone amides (6–8). This nest is formed by the N-terminus of an *α*-helix (9) and the preceding loop, a contiguous phosphate-binding fragment dubbed the Walker A motif (10). The Walker A motif is highly conserved across P-Loop NTPases and is characterized by the glycine-rich sequence GxxGxGK[T/S]. Guided by structural bioinformatics and laboratory-characterized model peptides, we and others have argued that these core features – a *βα* or *βαβ* motif bearing a glycine-rich phosphate binding loop – seeded the emergence of P-Loop NTPases (9, 11–15), as well as several other cofactor-associated domains, including the Rossmann (5, 16), HUP (17), and Nat/Ivy (18) evolutionary lineages.

The profound conservation and functional importance of the Walker A motif has led some to consider whether the motif may predate the *βα* element within which it is situated, hypothesizing that some aspect of structural and functional continuity within the P-Loop NTPase family extends back to a primordial peptide of just 6-8 resides in length (7, 8). The notion that folded domains may have evolved from short peptides is not without precedence: Dayhoff’s study of sequence duplications within ferredoxin (19) and Dutton’s subsequent demonstration of a metal cluster-supporting peptide maquette (20) have inspired research into the early evolution of cluster-associated protein folds (21) that, taken together, supports this view, albeit for a different class of enzymes.

In 2012, Bianchi and coworkers reported that the hexapeptide SGAGKT bound phosphate with microscopic *K*_D_ values ranging from about 10 µM to 1 mM (7). Although their methodology was unable to probe binding of backbone amides, the authors hypothesized that a stable peptide-phosphate complex could be indicative of a nest-like conformation reminiscent of contemporary enzymes (7, 8). This result – that a truncated, disembodied Walker A peptide retained significant affinity for phosphate – plus the hypothesized nest formation was an extreme example of functional continuity and would go on to shape thinking about the evolutionary trajectory of P-Loop NTPases for over a decade (13, 22–29). Yet, the relatively modest interaction energies accessible to phosphate binding, as opposed to the coordinate *covalent* Fe-S bonds of maquettes, would seem insufficient to drive structuring of a short glycine-rich peptide. Further, the significance of short *nucleotide*-binding peptides (30–33) – where interactions to the base may dominate – is unclear. Do Walker A-derived peptides have an intrinsic conformational preference that would support binding as nests? Are the interaction energies associated with nest binding modes sufficient to induce peptide structuring?

The question of whether a Walker A peptide sparked the emergence of P-Loop NTPases has significant implications for the evolution of anion-binding domains more generally, several of which host similar loop-nest structures (6). If true, (di)nucleotide binding domains would become quintessential examples of molecular ‘simple beginnings’ – complex structures that condensed around low complexity evolutionary nuclei. Thus, clarifying the extent to which a disembodied Walker A motif adopts a nest-like conformation upon binding to phosphate is fundamental to understanding the early evolutionary history of this fold and others. To this end, we have performed a detailed characterization of the dynamic properties of 7 representative Walker A motifs, including the previously-reported hexapeptide SGAGKT (7) and a number of control sequences, by Hamiltonian replica exchange (HREX) molecular dynamics (MD) to search for putative nest formation.

We find that the situated Walker A motif can be surprisingly independent, with some structures forming only a few interactions with the surrounding protein; and rigid, adopting a narrow distribution of conformations and undergoing only minor adjustments upon ligand binding. The disembodied motif, on the other hand, does not adopt nest-like conformations in MD simulations, though some intrinsic conformational preferences do exist. Uncorrelated preferences for nest-like dihedrals were observed, particularly of the glycine residues, but these preferences were modest. Inclusion of orthophosphate or GTP did not significantly induce nest formation, though GTP had a notably larger effect on the conformation of the peptide than orthophosphate, promoting bent peptide conformations. The GTP-bound bend conformations involved interactions with the entire nucleotide, not just the triphosphate group, suggesting that, from a biophysical standpoint, nucleotide binding may be a more accessible function than phosphate binding. Finally, a series of natural loops with the same amino acid composition but distinct evolutionary histories and situated-loop conformations (control loops) were found to behave comparably to Walker A peptides with respect to conformational dynamics and ligand binding.

Despite some flickers of nest-like conformations, we conclude that the Walker A motif – if it ever existed as a *free peptide in an aqueous environment* – is unlikely to have had significant nest-forming potential, even in the presence of phosphate or phospho-ligands. Moreover, the comparison to control loops indicates that a disembodied Walker A peptide is not a uniquely preferable solution to phosphate binding or nest formation under these conditions. Models of early P-Loop evolution must be revised. Finally, we argue that great care must be taken when interpreting the significance of highly conserved sequence fragments, which may represent eroded or incomplete molecular fossils.

## Materials and Methods

### Situated Loop Analysis

Representative P-loop NTPase structures were collected from the Evolutionary Classification of Domains (ECOD) database, version develop291 (34, 35). Domain sequences were clustered at 90% identity using CD-HIT (36), and structures with a resolution worse than 2.5 Å were excluded from further analysis. For each cluster, the liganded and unliganded structures with the highest resolution were taken to be the cluster representatives. Per residue distributions of dihedral angles and an analysis of supporting contacts were calculated using custom scripts (Zenodo DOI: 10.5281/zenodo.13685586) based on the PyMOL API (pymol.org) (37). Residues that form supporting interactions were identified by a heavy atom distance cutoff of 3.5 Å between the situated loop and the surrounding protein. Residues that are covalently attached to a loop residue, as well as the *i*+4 *α*-helix hydrogen bonding interactions at the C-terminal end of the of the situated loop, were excluded from the list of interacting residues as they are largely invariant.

### Selecting Walker A Sequences for Detailed Characterization

The P-Loop NTPase evolutionary lineage is associated with more than 120 distinct families (5). To identify families that are likely to have been present in the last universal common ancestor (LUCA), we calculated the phyletic distribution of each family across the microbial tree of life (**Table S1**). Hidden Markov Models (HMMs) for all known P-Loop NTPase families were downloaded from ECOD database (version develop279). (34, 35) These HMM profiles were then searched against the bacterial and archaea genomes in the curated Genome Taxonomy Database (38) (GTDB; release 95) using hmmsearch. A positive identification of a P-Loop NTPase domain, or hit, was defined as an independent expectation value (E-value) of ≤10^-4^ and an HMM profile coverage of ≥70%. The hmmsearch search space parameter, which is used in calculation of independent E-values, was set to 106,052,079 sequences (taken from Pfam (39)) to allow for facile comparison of E-values between runs. 15 families were present in ≥90% of phyla and ≥80% of species in both Archaea and Bacteria, of which 7 were chosen for detailed analysis (**Table S2)**. These seven families were chosen such that both common strand topologies within the P-Loop NTPase evolutionary lineage were represented. For each of these seven families, a representative structure with a resolution ≤2.0 Å was identified. Control sequences were chosen such that they have the same amino acid composition as one of the representative Walker A sequences but are from an unrelated protein lineage (**Table S3**).

### Starting conformations for the simulations

The simulation starting conformation for the hexapeptide SGAGKT (7) was generated by mutating the Walker A motif from G-protein p21^ras^, which has the sequence GGVGKS, in PyMOL (37) (ECOD domain e5p21A1; note that an ECOD domain identifier is constructed from the PDB ID (in this case 5p21 (40)), the chain identifier (A) and an ECOD domain identifier (1)). Starting conformations for the Walker A-derived and control octapeptides were taken from their respective crystal structures (**Table S2**). For simulations with GTP, the position of the ligand was either taken from the crystal structure directly or positioned by superimposition with a liganded structure. For simulations with HPO_4_^2−^, the position of the ligand was obtained by superimposition with the *β*-phosphate the previously positioned GTP.

### Hamiltonian Replica Exchange Molecular Dynamics Simulations

Hamiltonian replica exchange (HREX) molecular dynamics (MD) simulations (41) were performed on each Walker A-derived and control octapeptide in the presence and absence of the ligands orthophosphate or GTP. All Hamiltonian HREX-MD simulations simulations used the the Amber ff99SB-ILDN force field (42) and the TIP3P water model (43), as implemented in the GROMACS 2019.4 simulation package (44). The PLUMED v2.6. interface was used (45). Additional control simulations of the unliganded Walker A-derived octapeptides were performed using the CHARMM36m force field (46) to account for secondary structure bias of different force fields (47, 48). These simulations were prepared using the CHARMM-GUI (49). All ligand parameters are provided in the **Supporting Information**, **Tables S4** and **S5**. Following initial system preparation and equilibration (see **Supporting Information**), 1 µs HREX-MD simulations were performed on each system, with all solute atoms included in the hot region. This makes the HREX-MD simulations formally equivalent to REST2 simulations (41, 50). HREX-MD simulations were performed using a total of 8 replicas per system, with *λ* values scaled exponentially between 0.67-1. Exchanges were attempted every 4 ps, achieving an average exchange rate of ∼43% across all systems. Subsequent analyses, presented below, were performed only on the neutral replica (*λ*=1). Further details of the HREX-MD simulations are provided as **Supporting Information**.

### Simulation Analysis

Distance and angle analyses of the simulated peptides were performed using custom scripts (Zenodo DOI 10.5281/zenodo.13685586). Analysis of *φ*, *ψ* backbone dihedrals to detect *α*_R_ (R) and *α*_L_ (L) conformations of nest-forming residues was based on snapshots taken every 10 ps. The SGAGKT hexapeptide was aligned to the longer sequences shown in **Table S2** and renumbered accordingly for presentation purposes. Dihedral ranges were defined as −20°>*φ*>−140° and 40°>*ψ*>−90° for *α*_R_ and 20°<*φ*<140° and −40°<*ψ*<90° for *α*_L_ (6). For each frame, correlated nest-like conformations were assigned when consecutive residues adopt LR, RL, RLR, LRL, or LRLR conformations. Hydrogen bonds between the peptide and ligand were identified by calculating donor-acceptor distances and angles using GROMACS (44). Distance and angle cutoffs of 3.5 Å and 135° were used for the donor-acceptor distance and the donor-hydrogen-acceptor angle, respectively.

Dimensionality reduction was performed using time-lagged independent component analysis (tICA). Cosines of the *φ*, *ψ* backbone dihedrals along the entire peptide chain were used as input with a lag time of 1 ps. The resulting two-dimensional maps were then subject to K-Means clustering. Both the tICA analysis and K-Means clustering were performed using the MSMBuilder python package (51). Liganded and unliganded simulation for a given peptide data were concatenated prior to dimensionality reduction and clustering. For each trajectory, the root mean square deviation (RMSD) of the backbone heavy atoms with respect to the situated structure was calculated using the GROMACS internal RMSD function (44). Secondary structure analysis was performed on snapshots taken every 10 ps using MDTraj (52) and assigned using the Define Secondary Structure of Proteins (DSSP) nomenclature (53). For each residue, the percentage of time spent in each secondary structure conformation was calculated and averaged over the entire length of the peptide to obtain an estimate of the secondary structure content of the entire peptide.

## Results

### The Situated Walker A Loop Forms a Rigid, Pre-organized Nest for Phosphate Binding

The Walker A nest (**Figure 1A-B**) is situated between *β*1 and *α*1 of a P-loop NTPase domain. Nucleotide binding is dominated by backbone interactions, with up to 6 backbone amides forming hydrogen bonds to the *α* and *β* phosphate moieties of the ligand. The Walker A nest wraps around these phosphate moieties like a hand, rather unlike the phosphate binding loop of the Rossmann fold, which forms a comparatively flat binding surface. As binding occurs at the N-terminus of an *α*-helix, presumably the *α*-helix dipole contributes to binding as well (54). The primary side-chain interaction is a salt bridge between a phosphate moiety and the side chain of Lys7, though the details of this interaction depend on whether a diphosphate or triphosphate is bound. Many P-Loop NTPases also bind an Mg^2+^ dictation (**Figure 1B**, green spheres, **Table S6**), which is chelated by Ser/Thr8 and forms interactions with the phospho-ligand (but, critically, not the nucleophilic water molecule, *i.e.,* it is not involved directly in the chemical mechanism of catalysis (55)). Nucleotide binding, however, is not strictly Mg^2+^ dependent and, in structures without a dictation, the sidechain hydroxyl of Ser/Thr8 can interact directly with the ligand (as in e2gksA2 or e1k6mA1, see also **Table S6** for a distribution of contacts). The backbone dihedrals of residues 4-7 of the Walker A nest are characterized by an alternation between the right-handed *α*-helix (*α*_R_) and left-handed *α*-helix (*α*_L_) regions of the Ramachandran plot (**Figure 1C**). From residue 7 onward, the structure is a canonical right-handed *α*-helix. Unsurprisingly, the positions that adopt an *α*_L_ conformation are either exclusively (position 6) or mostly (position 4) Gly (**Figure 1D**).

**Fig. 1.**
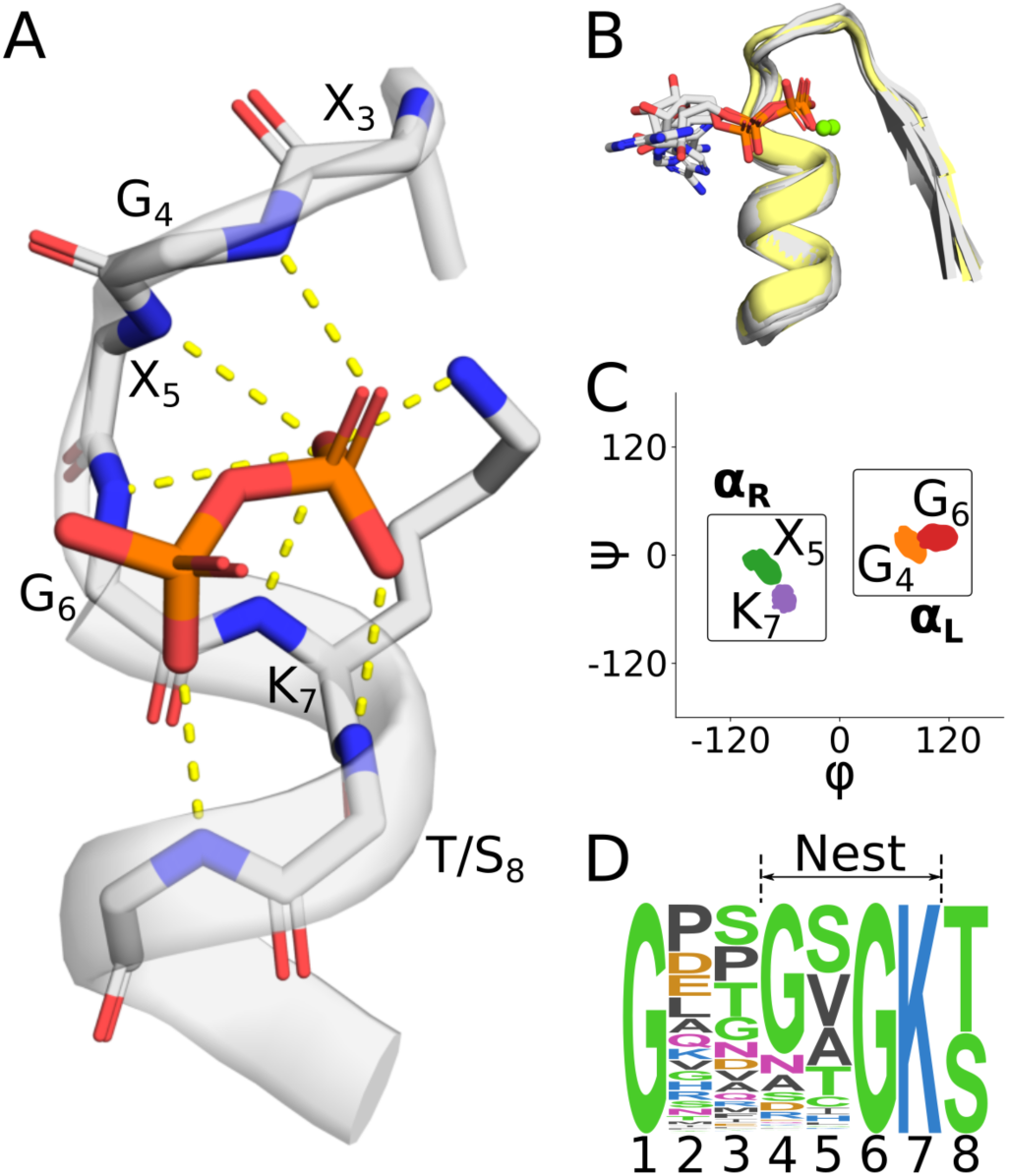
The phosphate binding nest of P-Loop NTPases. **A**. A representative phosphate binding nest, with 6 backbone amides and one lysine sidechain oriented towards and interacting (yellow lines) with the pyrophosphate moiety of ADP. ECOD domain e3k1jA2. **B**. Structural context of the Walker A motifs analyzed in this study. Structures with a bound phospho-ligand are colored gray; unbound structures are colored yellow. Mg^2+^ ions are drawn as green spheres and are present in only a subset of structures. Mg^2+^ ions were not included in our simulations, as described in the main text. ECOD domains: e3ievA2, e2it1A8, e3k1jA2, e5jrjA2, e6ojxA1, e2qy9A2. Note that the γ phosphate of the bound ATP was removed for clarity in e6ojxA1. **C**. Ramachandran plot showing the per-residue distribution of *φ*, *ψ* angles, including both liganded and unliganded structures. Boxes indicate the regions taken to be *ɑ*_R_ and *ɑ*_L_ in subsequent analyses. Note the alternating ɑ_L_-ɑ_R_-ɑ_L_-ɑ_R_ character of the nest. **D**. Sequence logo of the Walker A Motif demonstrating the canonical GxxGxGK[T/S] sequence pattern. P-Loop NTPase sequences were gathered from the PDB and filtered for 90% identity at the domain level.

Nucleotide binding does not dramatically alter the backbone conformation of the nest, in agreement with the observation that the phosphate binding nests of P-Loop NTPases tend to be relatively rigid (56–61). Both the structural overlay in **Figure 1B** and the Ramachandran plot in **Figures 1C** include liganded and unliganded structures, as well as structures with and without a bound dictation. A breakdown of *φ*, *ψ* angles for liganded and unliganded nests (**Figure S1**) shows that the dihedral distributions are overlapping for every residue of the putative nest. The largest adjustments relate to *φ*_6_, which has a difference between the means of 14 degrees. We conclude, then, that the backbone of the Walker A nest adopts a relatively rigid structure with complete or near-complete pre-organization for ligand binding. Stabilization of this conformation is likely due to a combination of intrinsic structural preference and supportive external interactions – the balance of which has implications for the potential adoption of nest-like structures by a disembodied Walker A peptide. An analysis of supportive interactions (**Figure S2**) revealed that while many Walker A motifs can make extensive contacts with the surrounding protein, there are also structures that make relatively few supportive interactions (2-15 contacts, on average 8; excluding backbone hydrogen bonds propagating the *α*-helix at the C-terminus and interactions to adjacent resides). Moreover, the specific supportive interactions themselves are not strictly conserved. Ultimately, we conclude that the Walker A structure is not likely to be encoded primarily through supportive interactions to interior residues of the motif.

### A Walker A-Derived Hexapeptide Does Not Form Nests

To better understand the conformational dynamics and evolutionary history of the Walker A motif, we analyzed the hexapeptide SGAGKT, which was previously hypothesized to form nests upon interaction with phosphate (6–8), by Hamiltonian replica exchange (HREX) molecular dynamics (MD) simulations. Simulations were carried out with and without ligands (HPO_4_^2−^ and GTP) for 1 μs and comprising 8 replicas with temperature range of 300-450 K per condition (see **Materials and Methods** for detailed information). We note here two important factors in our analysis: First, although primarily an ATP-binding sequence (10), both the Walker A and Walker B motifs occur in GTP-binding proteins, for example elongation factor Tu and Ras GTPase (62, 63). Indeed, initial studies of P-Loop NTPase nests focused on GTPases (7, 64). This apparent interchangeability stems from the fact that the situated Walker A peptide does not participate in base binding. Second, although phosphate binding of the Walker A motif typically involves metal dications, these metals are not required for ligand binding and are not a key determinant of nest formation (**Figure S3**). Consistent with the first experimental characterization of the Walker A-derived hexapeptide (7), we have not included metal cations in our analysis.

We find that the end-to-end distance (**Figure 2A**) and bend angle (**Figure 2B**) distributions are both broad, generally indicative of a flexible peptide. Nevertheless, a degree of conformational preference is apparent, with peaks corresponding to a tight hairpin (end-to-end distance of about 5 Å and a bend angle of about 45°). A time-lagged independent component analysis (tICA) of the MD trajectories (**Figure 2C**) reveals 6 primary clusters, the highest occupancy of which involves the peptide folding back onto itself, consistent with the end-to-end distance and bend angle distributions.

**Fig. 2.**
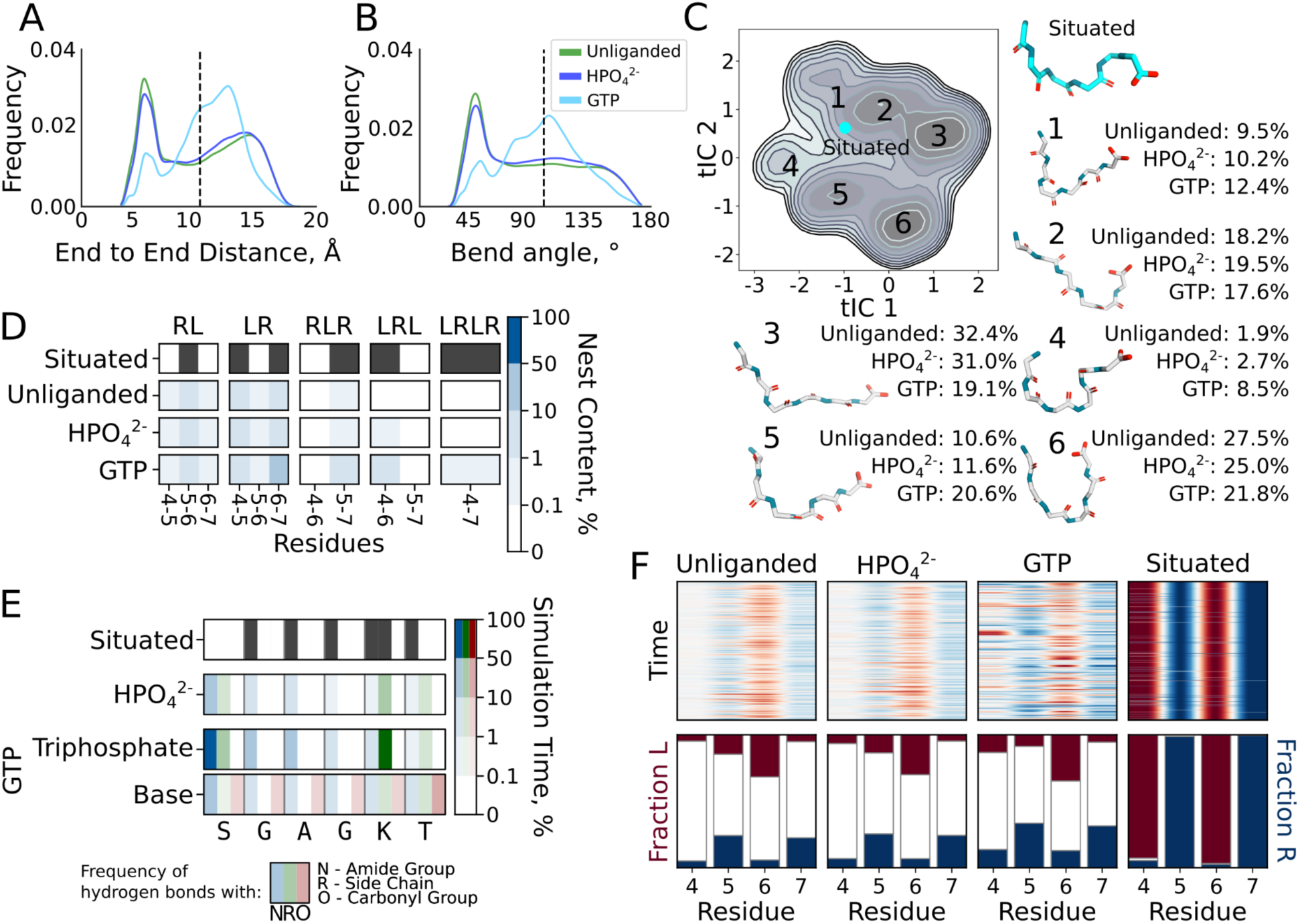
Conformational dynamics of the Walker A-derived hexapeptide SGAGKT. **A**. Distribution of C*_α_*_1_-C*_α_*_6_ distances for the free peptide in the presence and absence of a ligand. **B**. Distribution of C*_α_*_1_-N_4_-C*_α_*_6_ angles for the free peptide in the presence and absence of a ligand. The situated conformation is indicated with a dotted line in panels A and B. **C**. tICA projection derived from the dihedrals of all hexapeptide conformations, with and without ligand. The occupancy and a representative structure are provided for each of the 6 main clusters. The situated conformation is indicated by a cyan dot. **D**. Occurrence of correlated stretches of *α*_L_ and *α*_R_ conformations in each of our simulations, broken down per residue. **E**. Interaction profile of the peptide with a ligand. **F**. Uncorrelated preference of *α*_R_ or *α*_L_ backbone dihedrals in the presence and absence of phosphate ligands. The raw data for panels D and E is shown in **Tables S7** and **S8**, respectively.

Although the *global* similarity to the situated confirmation is low (**Figure 2A-C**), there is a modest preference for uncorrelated nest-like dihedral angles (**Figures 2F** and **S4**). Position 6 (Gly in the hexapeptide; motif numbering scheme in **Figure 1D**) has a greater propensity for *α*_L_ backbone dihedrals than do position 5 and 7, as in the situated conformation. In addition, positions 5 and 7 have a modest preference for the *α*_R_ backbone dihedral, once again reflecting the situated conformation. Correlated nest-like conformations between residues (**Figure 2D** and **Table S7**) are rare but not strictly absent from the simulations, with residues 5-7 having the greatest propensity to form nest-like regions, albeit with an occupancy of less than 1%.

Adding orthophosphate has a minimal effect on the conformation of the peptide (**Figure 2A-F**), with only a small increase in the nest-like conformation of residues 4-6. Addition of GTP yields a more pronounced effect – making the end-to-end distance, bend angle, and uncorrelated backbone dihedrals more *nest-like* but without inducing a global nest conformation (**Figure 2A-F**). The peptide-phosphate group interactions formed between the hexapeptide and either ligand (**Figure 2E**) are mediated largely by salt bridges between charged moieties (either the N-terminus or the side chain of Lys7) with a more modest contribution from the backbone amides, the hallmark feature of a nest binding mode. In contrast, a broad range of interactions involving the sugar and base of GTP were observed (**Figure S5** and **Tables S8** and **S9**). Thus, while flickers of nest-like conformations in the free Walker A peptide do occur, we do not observe global nest formation, nor do we detect persistent, organized phosphate binding mediated by the peptide backbone.

### Six Representative Octapeptides Do Not Form Nests

Six ubiquitously distributed P-Loop NTPase families were identified (**Table S2**). For each family, a representative structure was selected from the PDB (**Figure 1B**) and the conformational dynamics of its free Walker A motif analyzed as above. The selected Walker A octapeptides are identical at 3-7 of the 8 positions, owing to the diversity of the X positions, the occasional substitution of Gly4, and the use of Thr or Ser at position 8.

Like the hexapeptide, the octapeptides are highly dynamic but with flickers of nest-like conformations (**Figures 3A**-**B**, top row). As observed for the hexapeptide, the octapeptides have uncorrelated preferences for the situated backbone dihedral angles (**Figure 3A**) – with positions 5 and 7 generally adopting more *ɑ*_R_ dihedrals and positions 4 and 6 generally adopting more *ɑ*_L_ dihedrals. Surprisingly, even when position 4 is not Gly, a slight preference for *ɑ*_L_ can be retained (as in GKPNVGKS), though this is not always the case (as in GPESSGKT). The correlated nest propensities (**Figure 3B; Tble S7**) show some notable differences between the peptides: First, a greater propensity for correlated nest-like structures are observed for almost all of the octapeptides relative to the hexapeptide. Second, clear differences are observed between the octapeptides, suggesting that the identity of the X residues also augments the nest forming potential. We note that the peptide with the greatest correlated nest-like conformations (GEPGTGKS) has a proline before Gly4, which may play a key role in restricting the conformational space of the peptide and promoting nest-like conformations that involve three residues (residues 4-6, 7.1% occupancy; compared to 0.5-1.7% occupancy for all other octapeptides).

**Figure 3.**
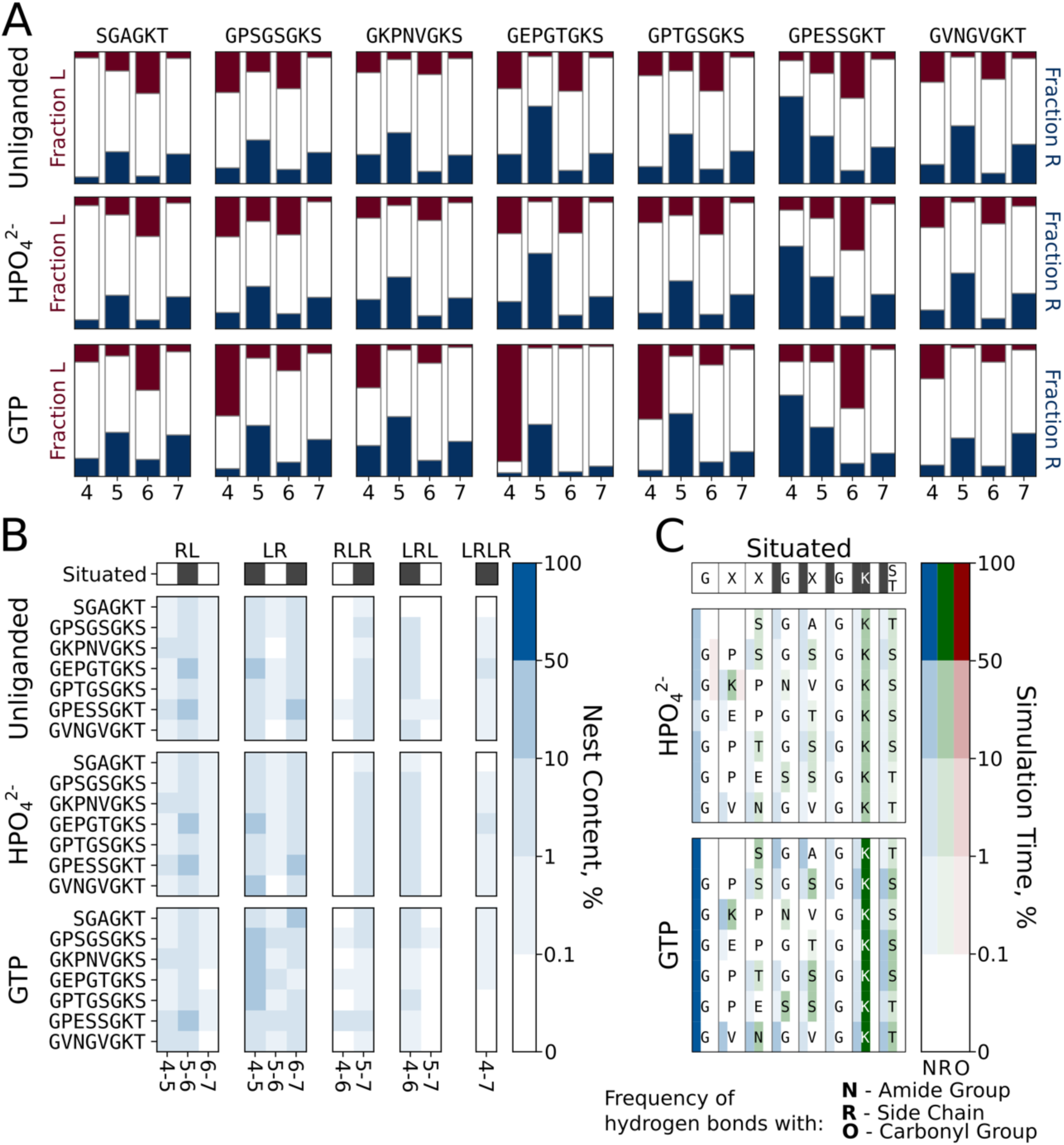
Conformational dynamics of Walker A-derived octapeptides. **A**. Uncorrelated preference for *α*_L_ and *α*_R_ backbone dihedrals. **B**. Occurrence of correlated stretches of *α*_L_ and *α*_R_ conformations. **C**. Interaction profile of the peptides with a ligand. Plot for contacts with GTP shows cumulative probability for interaction with all 3 phosphates groups of GTP. The raw data for panels B and C is shown in **Tables S7** and **S8.**

The inclusion of phosphate has only a minor effect on the properties of the octapeptides (**Figures 3 A-B**, middle row; **Table S7**), consistent with the hexapeptide results. GTP (**Figures 3 A-B**, bottom row; **Table S7**) inclusion had a more dramatic effect, increasing nest-like dihedrals at positions 4-5 by ∼15-30% occupancy for three octapeptides. This increase in nest character, however, does not translate to longer stretches of nest-like residues. As before, (tri)phosphate binding is mediated largely by interactions involving the charged N-terminus and Lys7, with a comparatively modest contribution from backbone amides (**Figure 3C**, **Table S8**). Interactions with the GTP sugar and base are more varied – with participation of the N-terminus, backbone carbonyls, polar/charged side chains, and backbone amides (**Figure S5**) – and GTP binding was significantly more persistent than binding of orthophosphate (>99% vs ∼30% simulation time, respectively; **Table S9**). In short, robust adoption of nest-like conformations was not observed for the octapeptides, even in the presence of phospho-ligands. Simulations performed with a different force field echo these results (**Figure S6, Table S10**). Even adding a distance restraint that penalizes deviation of the N- and C-termini from the end-to-end distance of the situated motif (a crude mimic of the effect of the protein scaffold on P-loop structure) did not induce global nest formation or nest-like binding, though it did increase the partial nest character of the peptides (**Figure S7, Tables S11** and **S12**).

### Walker A Peptides Behave Similarly to Control Peptides

Six control loops were selected that have the same amino acid composition as a Walker A octapeptide but different sequences and situated conformations (**Figures 4A** and **B, Table S3**). In most cases, the control loops are not associated with phosphate binding or a highly conserved sequence motif, and are thus unlikely to have undergone an evolutionary trajectory from a short peptide to a folded domain, as has been proposed for P-Loop NTPases. Analyzing the properties of these loops, then, can serve as a useful point of comparison with which to judge the properties of the octapeptides. Foremost, and particularly in the absence of ligand (**Figure 4**) or with orthophosphate (**Figure S8**), the control loops and the Walker A octapeptides have essentially equivalent dynamic properties: Broad distributions of end-to-end distances (**Figure 4C**) and bend angles (**Figure 4D**), though often with some conformational preference, and a similar degree of structural similarity between the situated and the free peptide (**Figure 4E**). Although the patterns of correlated and uncorrelated nest-like conformations (**Figure S9**) were different from the Walker A peptides, their magnitude was roughly the same – suggesting that this degree of structuring is common among free peptides. As with the other loops analyzed, the control loops did not bind persistently to orthophosphate (25.2-53.0% of simulation time; slightly higher than for the Walker A octapeptides, which ranged from 24.7-39.9% of simulation times) and interacted predominantly via charged groups and the N-terminus of the peptide (despite the different locations of these residues, **Figure S9**). Both the Walker A octapeptides and the control peptides adopt similar secondary structure distributions with or without added ligands (**Figure S10**).

**Figure 4.**
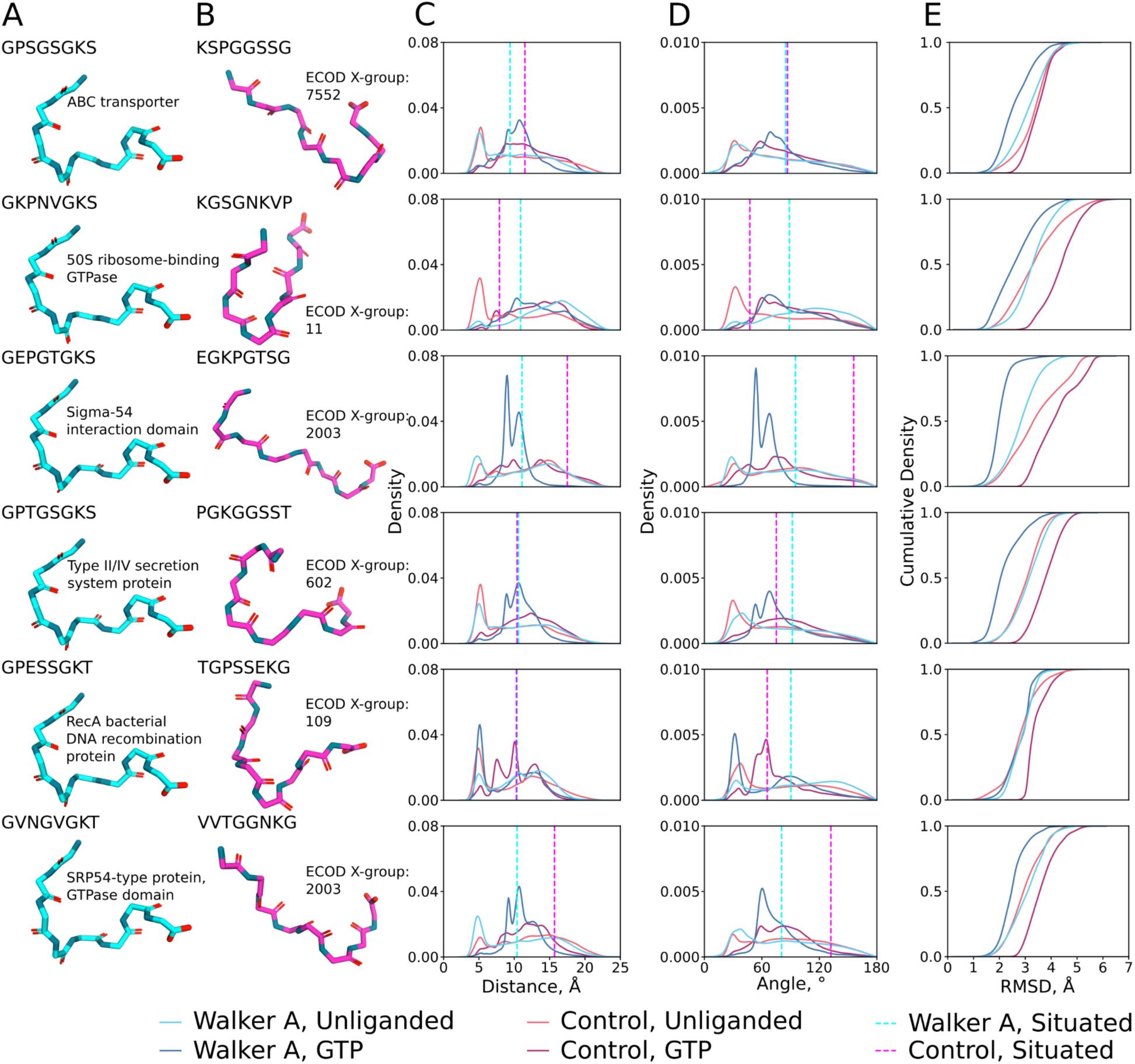
Conformational dynamics of the control octapeptides. **A**. Situated structures of the Walker A derived octapeptides. **B**. Situated structures of control octapeptides. **C**. Distribution of C*_α_*_1_ - C*_α_*_8_ distances for the free peptide in the presence and absence of GTP. **D**. Distribution of C*_α_*_1_-N_5_-C*_α_*_8_ angles for the free peptide in the presence and absence of GTP. The situated conformation for panels C and D is indicated with a dotted line. **E**. Cumulative distribution of the root mean square deviations (RMSD) of each set of simulations towards the situated structure of the respective peptide in the presence and absence of GTP.

Once again, interactions with GTP caused larger shifts in the conformations of the peptides than orthophosphate, no doubt due to its comparatively large interaction surface (**Figure 4**). Intriguingly, whereas the Walker A peptides generally become more similar to their situated structure (**Figure 4D**), the control peptides become less similar to their situated structures, despite having similar overall interactions with different parts of the nucleotide (**Table S9**, although note that some peptide-base interactions are stronger in the Walker A than in the control sequences, see **Figure S5**). This result is likely because GTP induces a bend in the peptide that is similar to the bend angle of the situated Walker A loop but rather dissimilar to the bend angles of several situated control loops (**Figures 4A** and **B**).

## Discussion

The limits of evolutionary continuity, particularly of P-Loop NTPases, is a topic of intense interest (5, 7–9, 11, 13–15, 65, 66). There is a broad consensus that the origin of this family lies somewhere between a short Walker A peptide that forms a nest upon phospho-ligand binding (7, 8) and a short *βα* or *βαβ* peptide that can either fold independently as a monomer or upon oligomerization (5, 11, 13–15). Whereas the potential for fold evolution from an oligomerizing peptide composed of several secondary structure elements has been demonstrated for multiple folds (13, 67–70) – making this a viable degree of complexity for an evolutionary starting point – evolutionary continuity down to a simple, short peptide of ∼10 residues is less well established, with FeS maquettes being the primary example (20, 71). Nevertheless, short nucleotide-binding peptides have been reported (30–33), speaking to the chemical plausibility of this evolutionary scenario for phospho-ligands. The present evolutionary scenario depends on two key points: (1) The disembodied Walker A motif binds phosphate or a phospho-ligand and (2) the binding mode of the Walker A-derived peptide is nest-like in structure. These two points, which concern different aspects of continuity -- one related to function and the other to structure -- will be discussed in turn.

Initial studies on the hexapeptide observe binding of orthophosphate using potentiometric titrations (7) that are sensitive to shifts in p*K*_a_, and thus cannot probe binding by backbone amides. These results are in rough agreement with the MD simulations, where orthophosphate binding was observed in roughly 30% of frames and was mediately largely by charged groups. However, the Walker A octapeptides and the control peptides were highly similar with respect to both binding patterns and binding persistence, suggesting that interactions observed by potentiometric titration may have been largely non-specific. Although binding to nucleotides was more persistent (>99% of simulation time in all cases) and induced larger structural changes, the Walker A peptides and the control peptides once again exhibited similar properties. Taken together, we conclude that a disembodied Walker A peptide is unlikely to be a privileged phospho-ligand binder; instead, a diversity of peptides with equivalent or superior binding properties likely exist.

The sustained GTP binding (**Table S9**) in our simulations may be noteworthy, given that a variety of short peptides bind nucleotides (usually ATP), including peptides using prebiotic amino acids (31–33). One caveat to this observation is that these peptides were typically engineered with sequences specifically selected for their ability to bind ATP (31). The short ATP-binding octapeptide TACGQKSP, on the other hand, was extracted from the ATP binding site of frog actomyosin and binds MgATP with a *K*_D_ of 0.22 mM (30). Similar to our GTP simulations (**Figure S5** and **Tables S8** and **S9**), this peptide binds ATP *via* interactions with all parts of the ligand, not just the triphosphate, and discriminated between ATP, GTP, and CTP binding, with ATP binding preferred (30). These data suggest that it is possible to achieve ligand binding from short-disembodied peptides, particularly of nucleotides where supportive interactions with the base can be formed. Indeed, peptide-base interactions could have played an important role in substrate discrimination in an ATP-rich prebiotic world (72).

While some aspect of functional continuity seems at least plausible, the question of structural continuity is unambiguous: Free Walker A-derived peptides do not have a significant preference for nest-like conformations, even in the presence of phospho-ligands. The current data suggest that the Walker A motif and the *βαβ* peptide within which it was situated co-evolved – with the loop and the surrounding peptide reciprocally adjusting until the canonical binding mode was found. Although flickers of nest-like conformations are observed, these are weak and more reasonably interpreted as residual preference for nest-like conformations imparted by evolution operating on the situated peptide. The high content of Gly within the peptide may be a signature of this co-evolution, as high Gly content in a free peptide is expected to disfavor folding and collapse (though it must also be noted that Gly is the simplest amino acid and may have enjoyed high prebiotic availability). If a strong propensity for nest formation had been observed, then the framing of this structural element as an evolutionary nucleus would, in our view, have been supported.

The field of protein evolution continues to grapple with how to interpret profound sequence conservation – is it evidence of an evolutionary nucleus, the earliest traces of a fold contained with a short peptide? Or, are these signals fragmentary – motifs that resisted erosion over time but nonetheless existed within a specific context that is no longer recoverable, and thus not the “full” evolutionary seed. Disentangling these scenarios is difficult. Lupas’ work on searching for a “primordial peptide vocabulary” has sought to account for this, in part, by finding conserved sequence-structure elements within multiple, distantly related contexts (11). Likewise, Kolodny *et al*., proposed the concept of “bridging themes,” which takes a related approach, except that structure similarity is not required, allowing the inclusion of metamorphic fragments as well (73). The latter approach was instrumental in uncovering the potential deep evolutionary connection between P-Loop NTPases and Rossmann enzymes (73). Although reasonable, these approaches are imperfect, and the identified fragments may have emerged within a fold, as an assembling oligomer, or even folded independently. As the interfaces of the conserved fragments were overwritten to adapt to changing contexts, information about the originating context was lost or distorted. For the Walker A-derived peptides, the presence of Mg^2+^ or the ability to assemble may reveal a yet-unappreciated aspect of evolutionary continuity between a Walker A peptide and the NTPase evolutionary lineage. It is only through extensive characterization that the relative likelihood these scenarios can be assessed. At present, the best evidence for the evolutionary seed of P-Loop NTPases pointes to oligomerizing *βαβ* or *βα* peptides.(13)

## Acknowledgements

This work was supported by the Knut and Alice Wallenberg Foundation (grant numbers 2018.0140 and 2019.0431). The simulations were enabled by resources provided by the National Academic Infrastructure for Supercomputing in Sweden (NAISS), partially funded by the Swedish Research Council through grant agreement no. 2022-06725. P.L. was supported by the Okinawa Institute of Science and Technology Graduate University (OIST) with subsidy funding from the Cabinet Office, Government of Japan. We also want to thank Dan Tawfik for inspiring this work and for helpful discussion. L.M.L. is grateful to Eric Smith for insightful discussions.

