## Supporting Information for "Redefining the Limits of Functional Continuity in the Early Evolution of P-Loop NTPases"

**This PDF file includes:**

Supporting text
Figures S1 to S10
Tables S1 to S15
SI References

**Supporting Information Text**

**Supplementary Methods - System Preparation and Equilibration Procedure for Hamiltonian Replica Exchange Molecular Dynamics Simulations**

All systems were placed in triclinic water boxes of TIP3P (1) water molecules, with box lengths defined so that all atoms were at least 10 Å from the box edges. All simulations were performed using periodic boundary conditions, with long-range electrostatics being handled using the Particle Mesh Ewald (PME) approach (2), and a 10 Å cutoff. $K^+$ and $Cl^-$ counterions were added to each system to neutralize the charge of each peptide/complex, and to give a final ion concentration of 0.1 M. All simulations were performed using a 2 fs time step, with all bonds containing hydrogen atoms being constrained using the SHAKE algorithm (3).

Each system was first minimized using 500 steps of unrestrained steepest descent minimization, followed by 5000 steps of conjugate gradient minimization with constraints on all bonds with hydrogen atoms. At this point, the system was then heated from 0 to 300K over 400 ps of simulation time in an NVT ensemble, with temperature and pressure regulated using velocity rescaling (4) (collision frequency of 1 ps). Following this, a further 400 ps of equilibration was performed at 300K in an NPT ensemble, once again using velocity rescaling and the Parrinello-Rahman barostat (5), with 1000 kJ $mol^{-1}$ $nm^{-1}$ harmonic restraints applied to all heavy atoms of the peptide. Both NVT and NPT equilibration simulations were performed using a time step of 2 fs and the P-LINCS (6) algorithm for restraining all bonds to hydrogen atoms.

Finally, in the case of the end-restrained HREX-MD simulations, an additional 4184 kJ $mol^{-1}$ $nm^{-1}$ harmonic restraint was put on the $C_{\alpha1}$ - $C_{\alpha8}$ distance corresponding to that distance in the respective situated structure (see **Table S15**).

**Supplementary Figures**

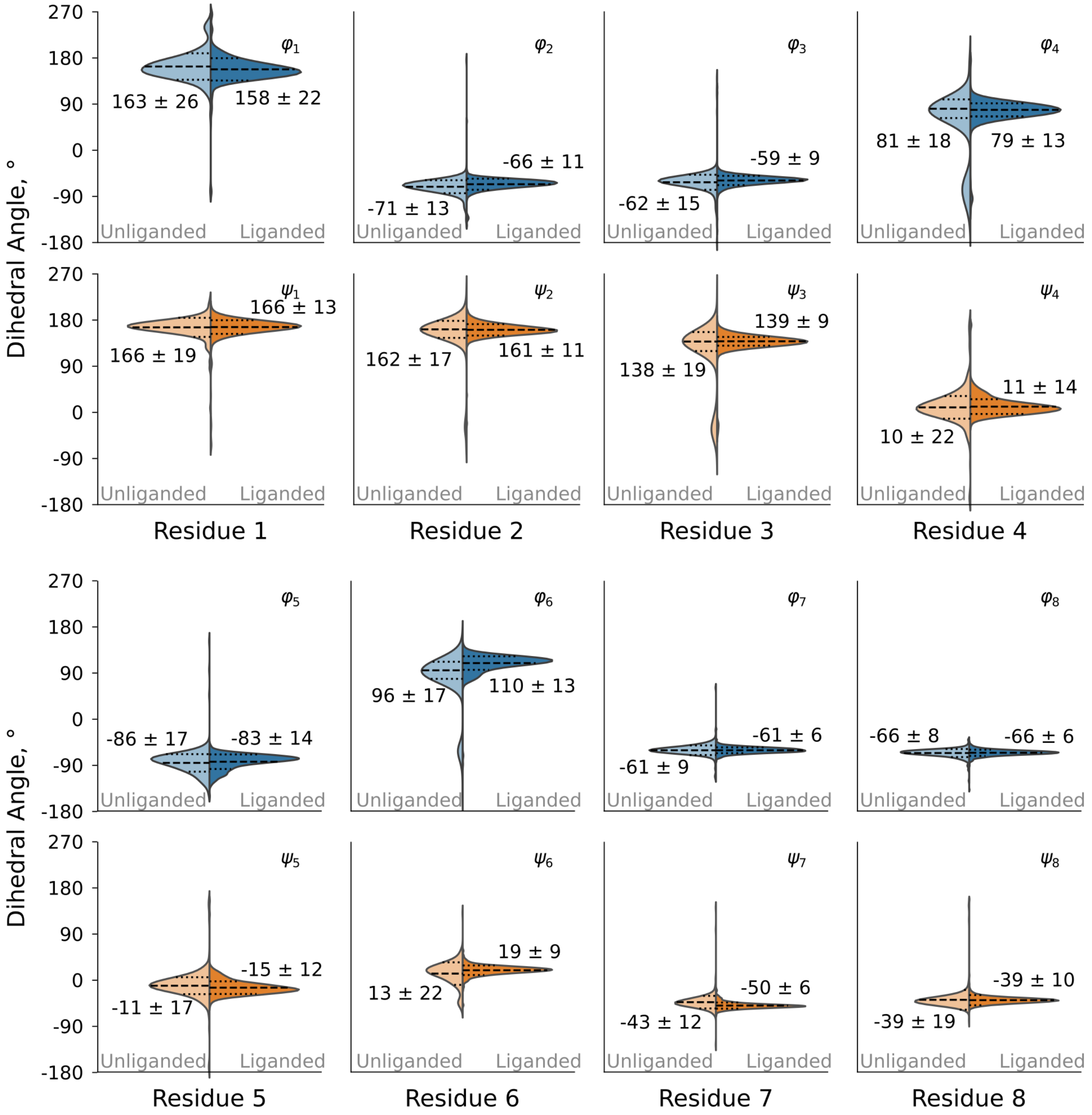


**Fig. S1.** Analysis of Walker A P-Loop conformational change upon phospho-ligand binding. Shown here are the distributions of the *φ*- and *ψ*-dihedral angles for representative liganded or unliganded domains. The mean values for each distribution are shown using dashed lines and the standard deviation ranges are shown using dotted lines.

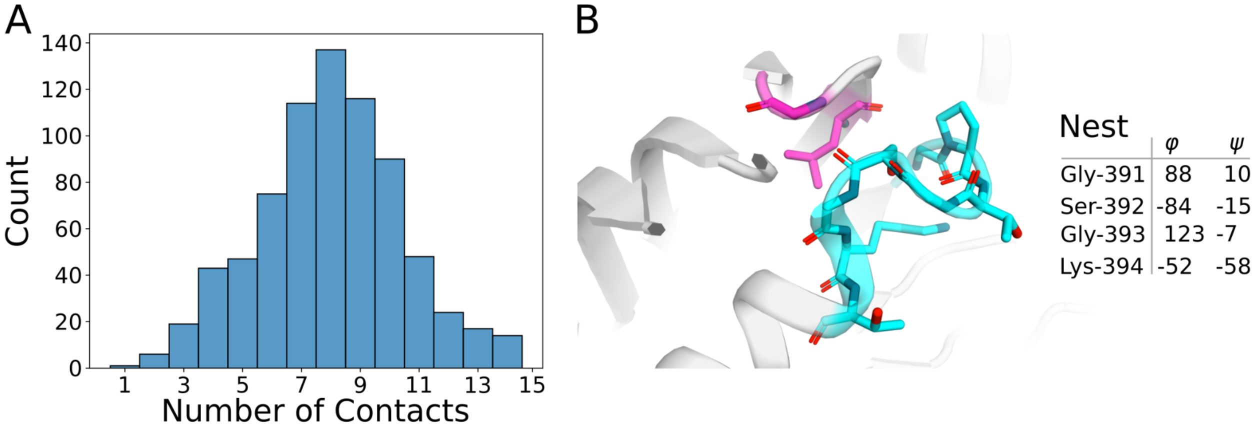


**Fig. S2.** Supporting interaction analysis. **A.** Distribution of the number of heavy atom contacts formed by situated Walker A motifs with surrounding protein structure. *α*-helical contacts, which are formed by the C-terminal residues of the situated Walker A motif, and interactions to covalently connected, adjacent residues were excluded from the analysis as they are largely invariant. The cutoff distance used for calculating contacts was 3.5 Å. **B**. Example of a P-loop with a canonical conformation and a small number of supporting contacts (ECOD domain e4q7mB1, with two supporting contacts). The P-loop is shown in cyan and residues within 3.5 Å that are not related to propagation of the *α*-helix are shown in magenta. Backbone *φ*- and *ψ*-dihedral angles of the nest-forming residues in this structure are shown in the table.

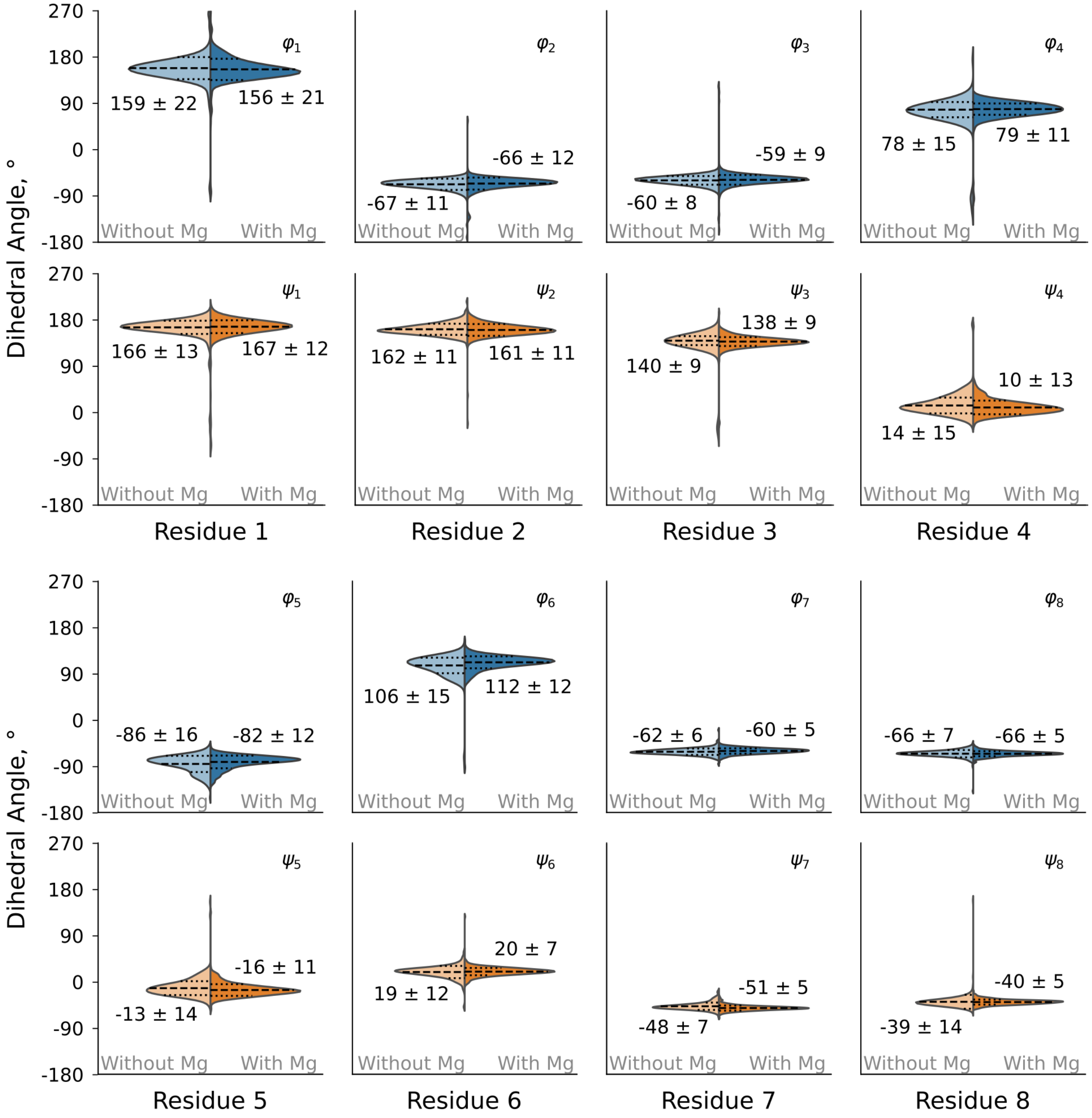


**Fig. S3:** Analysis of Walker A P-Loop conformational change upon $Mg^{2+}$ binding. Shown here are the distributions of the $\varphi$- and $\psi$-dihedral angles for representative structures with and without a bound $Mg^{2+}$. The mean values for each distribution are shown using dashed lines and the standard deviation ranges are shown using dotted lines.

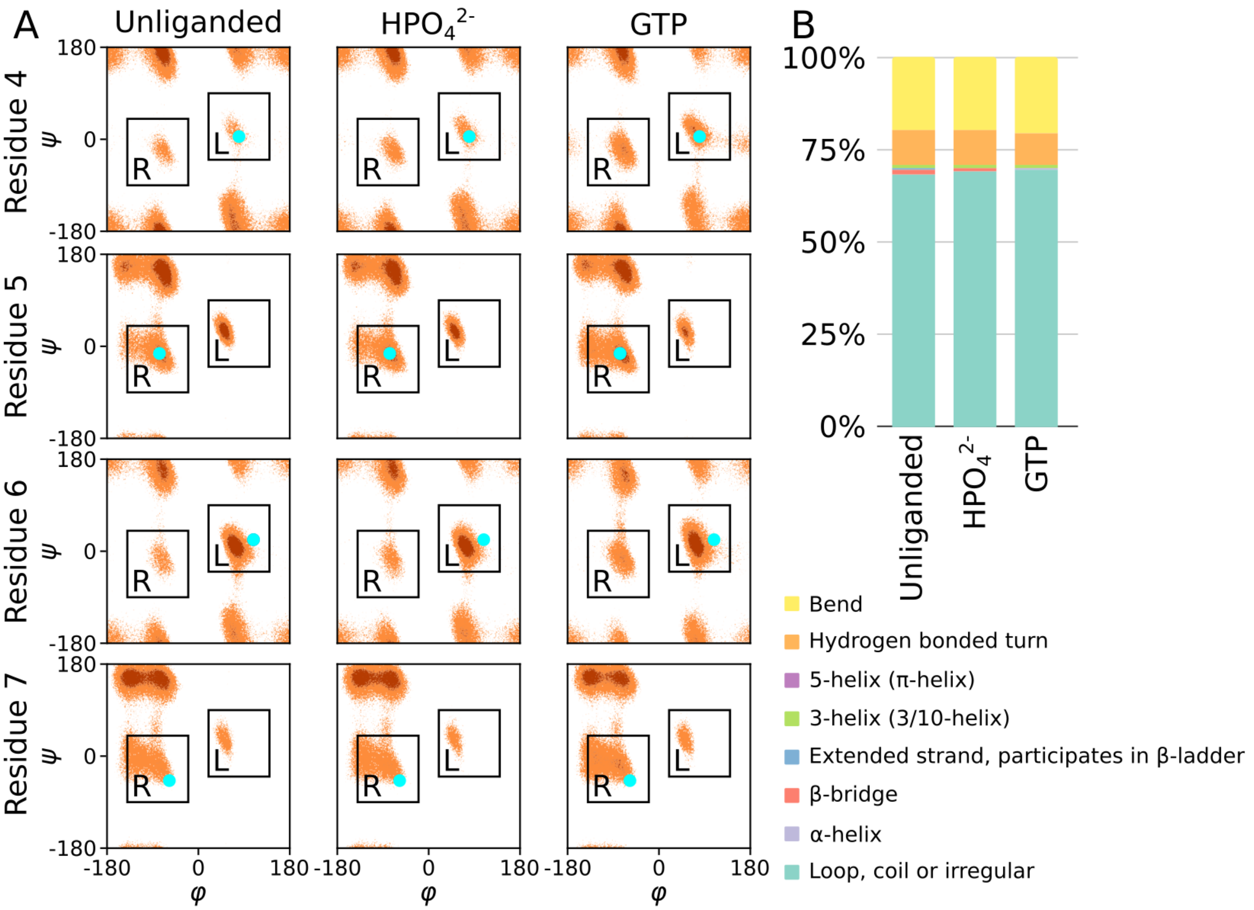


**Fig. S4.** Conformational dynamics of the Walker A-derived hexapeptide SGAGKT (see ref. (7)) from 1 µs HREX simulations in the presence ($HPO_4^{2-}$ or GTP) or absence of a ligand. **A**. Calculated Ramachandran plots. The values for the situated structures are indicated with a cyan dot. **B**. Secondary structure composition calculated using MDTraj (8) and presented using DSSP (9) annotations.

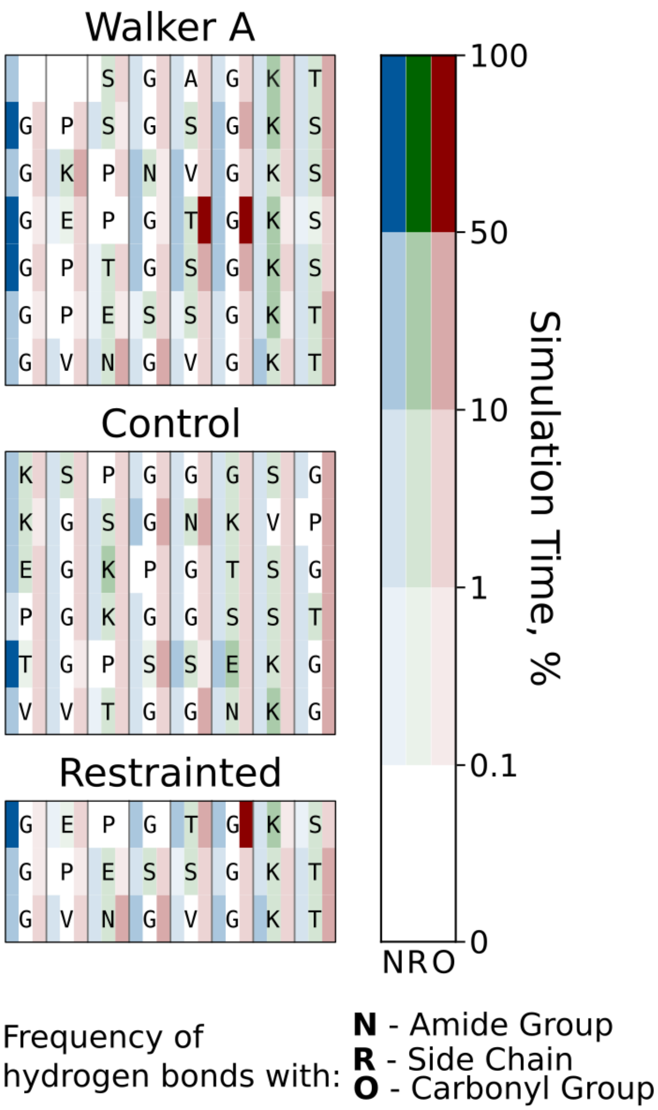


**Fig. S5**. Interaction profile of the Walker A peptides with the base and sugar groups of GTP. Hydrogen bond donor-acceptor distance and angle cutoffs were 3.5 Å and 135°, respectively. The raw data for this figure is shown in **Tables S8, S12** and **S14.**

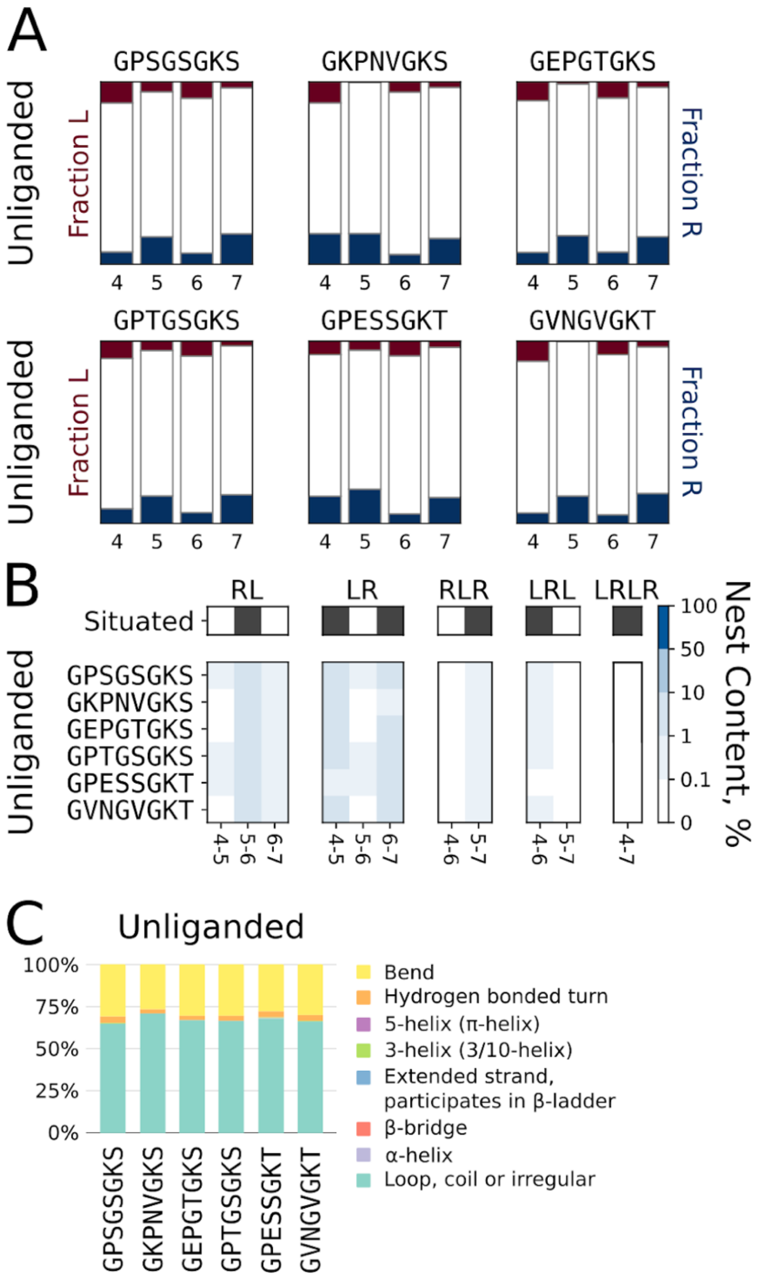


**Fig. S6.** Conformational dynamics of the Walker A-derived octapeptides during simulations using the CHARMM36m force field.(10) **A.** Uncorrelated preferences for $\alpha_L$ and $\alpha_R$ backbone dihedrals. **B.** Occurrence of correlated stretches of $\alpha_L$ and $\alpha_R$ conformations. **C**. Secondary structure calculated by MDTraj (8) and presented using DSSP (9) annotations. The raw data for panel B is shown in **Table S10.**

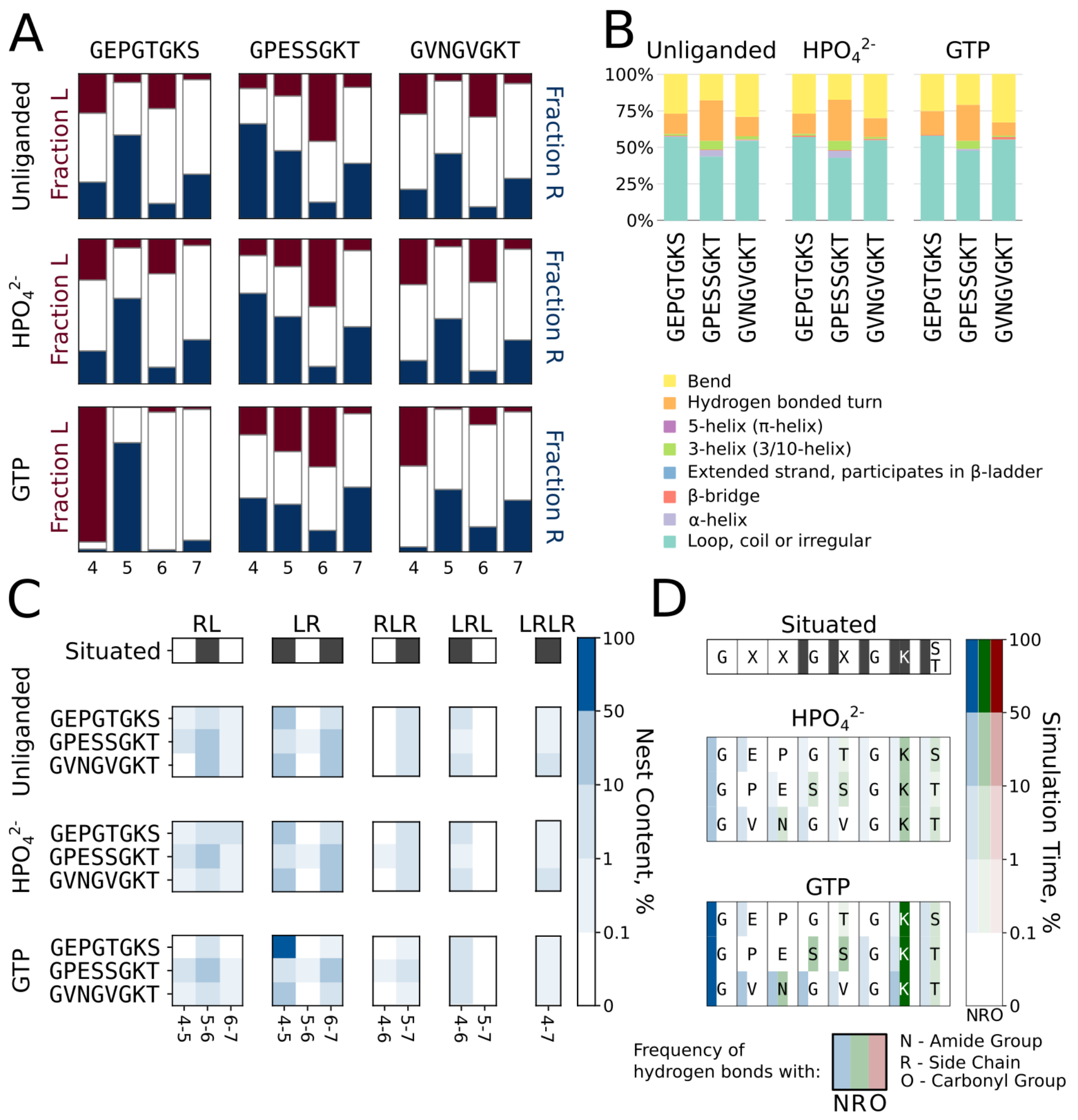


**Fig. S7.** Conformational dynamics of the Walker A-derived octapeptides in simulations in which the $C_{\alpha 1}$-$C_{\alpha 8}$ distance of the peptide is restrained. **A.** Uncorrelated preferences for $\alpha_L$ and $\alpha_R$ backbone dihedral angles. Secondary structure calculated by MDTraj (8) and presented using DSSP (9) annotations. **C**. Occurrence of correlated stretches of $\alpha_L$ and $\alpha_R$ conformations. **D**. Interaction profile of the peptides with a ligand. Hydrogen bond donor-acceptor distance and angle cutoffs were 3.5 Å and 135°, respectively. The raw data for panels C and D is shown in **Tables S11** and **S12.**

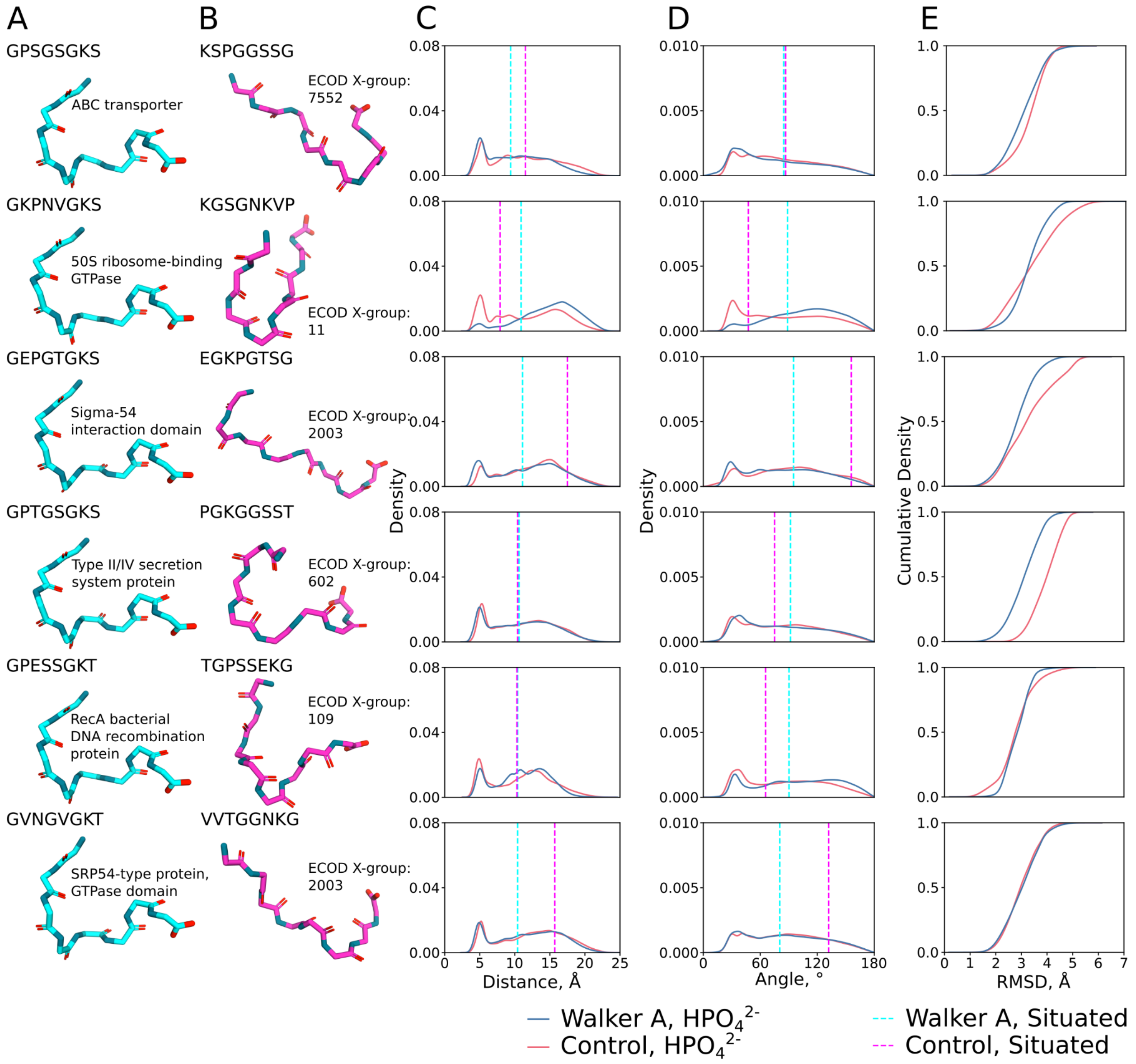


**Fig. S8.** Conformational dynamics of the control octapeptides in presence of $HPO_4^{2-}$. **A**. Situated structures of the Walker A-derived octapeptides. **B**. Situated structures of control octapeptides. **C**. Distribution of $C_{\alpha1}$ - $C_{\alpha8}$ distances for the free peptide in the presence of a phosphate ligand. **D**. Distribution of $C_{\alpha1}$-$N_5$-$C_{\alpha8}$ angles for the free peptide in the presence of a phosphate ligand. The situated conformation for panels C and D is indicated with a dotted line. **E**. Cumulative distribution of the root mean square deviations (RMSD, Å) of each set of simulations in the presence of orthophosphate relative to the situated structure.

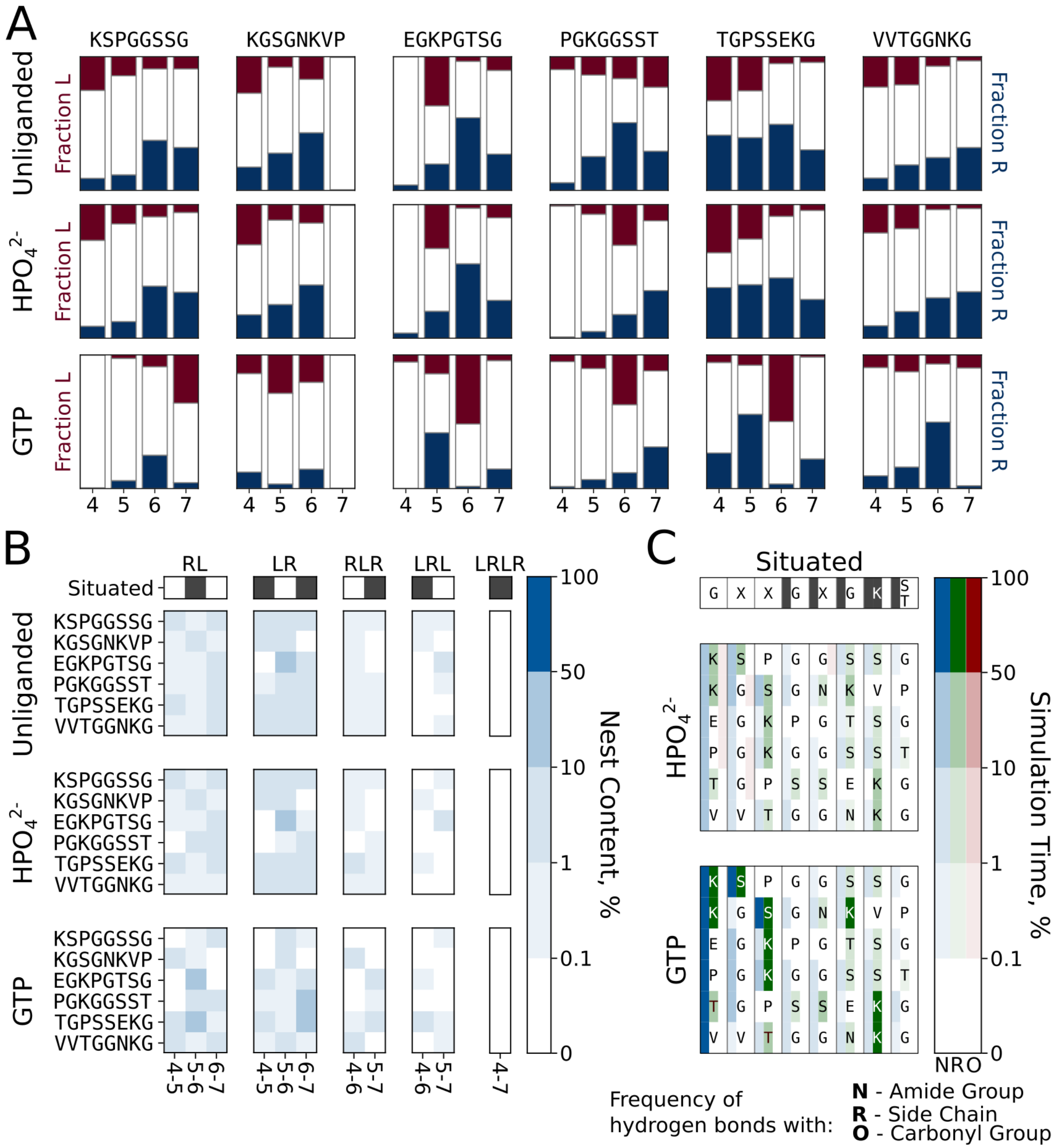


**Fig. S9**. Conformational dynamics of control octapeptides. **A**. Uncorrelated preference for $\alpha_L$ and $\alpha_R$ backbone dihedrals. **B**. Occurrence of correlated stretches of $\alpha_L$ and $\alpha_R$ conformations. **F**. Interaction profile of the peptides with a ligand. The raw data for panels B and C is shown in **Tables S13** and **S14.**

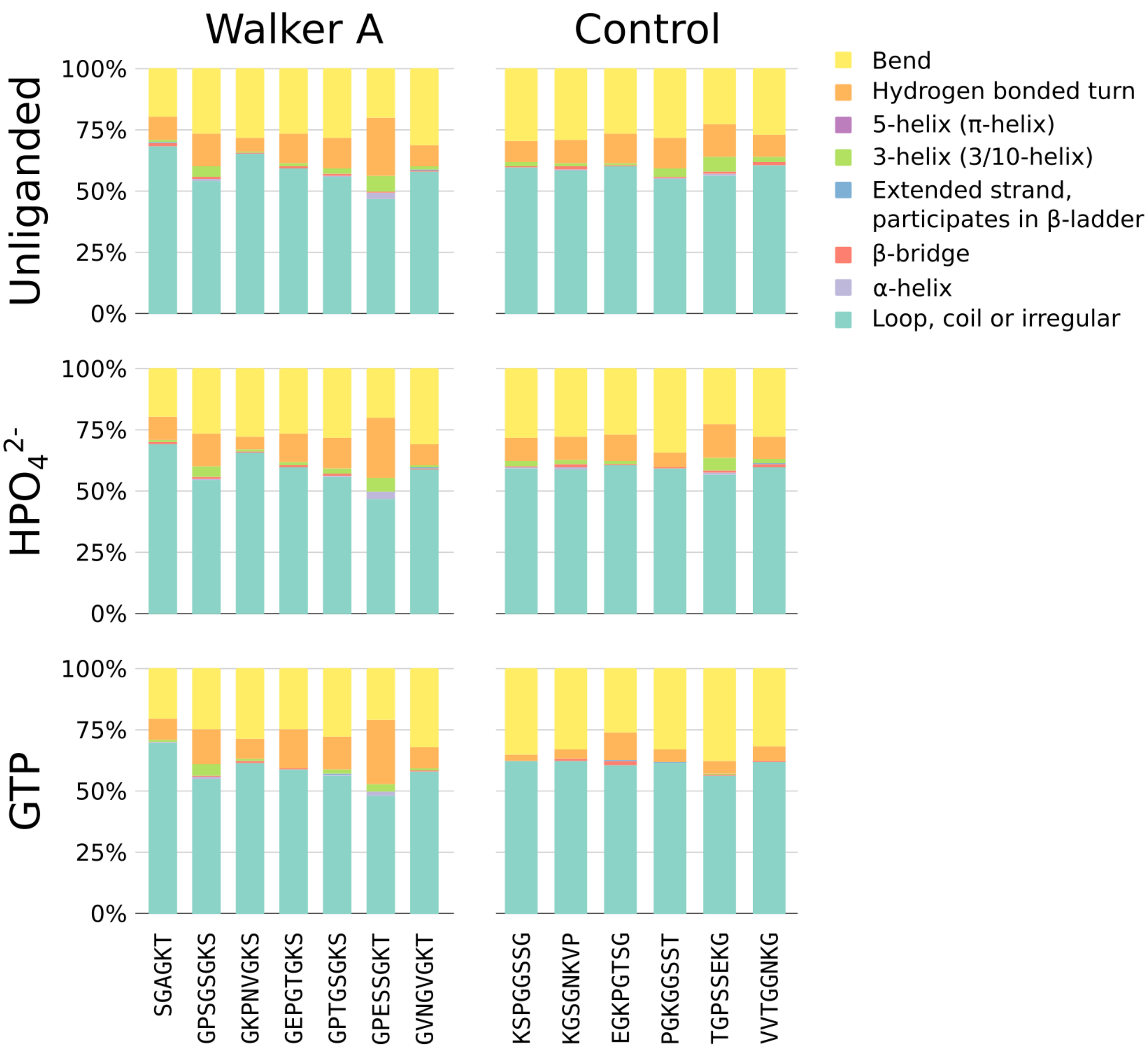


**Fig. S10**. Secondary structure compositions for the disembodied Walker A peptides and the control peptides in the presence ($HPO_4^{2-}$ or GTP) or absence of a ligand. Calculated using MDTraj (8) and presented using DSSP (9) annotations.

### Supplementary Tables

**Table S1.** Phyletic distribution of P-loop NTPase families across the microbial tree of life.[a]

| F-Group Name | Frac. Species, Archaeal | Frac. Species, Bacterial | Frac. Phyla, Archaeal | Frac. Phyla, Bacterial |
|---|---|---|---|---|
| AAA_31 | 0.95 | 0.96 | 1.00 | 1.00 |
| ABC_tran | 1.00 | 1.00 | 1.00 | 1.00 |
| ABC_tran_1 | 1.00 | 1.00 | 1.00 | 1.00 |
| DEAD | 1.00 | 1.00 | 1.00 | 1.00 |
| DEAD_1 | 1.00 | 1.00 | 1.00 | 1.00 |
| MnmE_helical_2nd | 1.00 | 1.00 | 1.00 | 1.00 |
| Sigma54_activat | 1.00 | 1.00 | 1.00 | 1.00 |
| AAA_31_1 | 0.93 | 0.95 | 1.00 | 0.99 |
| DEAD_3 | 1.00 | 0.99 | 1.00 | 0.99 |
| T2SSE | 0.93 | 0.80 | 1.00 | 0.99 |
| SMC_N | 0.77 | 0.96 | 1.00 | 0.98 |
| RecA | 0.93 | 0.93 | 1.00 | 0.98 |
| Helicase_C_1 | 0.97 | 0.96 | 1.00 | 0.96 |
| CTP_synth_N | 0.85 | 0.90 | 1.00 | 0.95 |
| SRP54 | 0.97 | 0.93 | 1.00 | 0.91 |
| Helicase_C | 0.53 | 0.51 | 1.00 | 0.83 |
| Beta-Casp | 0.96 | 0.34 | 1.00 | 0.70 |
| DUF853 | 0.69 | 0.36 | 1.00 | 0.65 |
| AAA | 0.96 | 0.07 | 1.00 | 0.48 |
| Ras_1 | 0.75 | 0.99 | 0.94 | 0.98 |
| Thymidylate_kin | 0.89 | 0.88 | 0.94 | 0.98 |
| SMC_N_1 | 0.57 | 0.68 | 0.94 | 0.88 |
| Helicase_C_2 | 0.82 | 0.53 | 0.94 | 0.77 |
| DEAD_2 | 0.74 | 0.46 | 0.94 | 0.76 |
| Cytidylate_kin2 | 0.75 | 0.21 | 0.94 | 0.65 |
| Helicase_C_5 | 0.85 | 0.28 | 0.94 | 0.58 |
| MCM_AAA | 0.69 | 0.01 | 0.94 | 0.27 |
| POR | 0.88 | 0.64 | 0.89 | 0.98 |
| Ras | 0.67 | 0.98 | 0.89 | 0.97 |
| ATP-synt_ab | 0.83 | 0.97 | 0.83 | 1.00 |

| | | | | |
|---|---|---|---|---|
| CobA_CobO_BtuR | 0.41 | 0.43 | 0.83 | 0.79 |
| sufC | 0.55 | 0.64 | 0.83 | 0.60 |
| SKI | 0.36 | 0.89 | 0.78 | 0.93 |
| UvrD_C | 0.36 | 0.97 | 0.72 | 0.98 |
| SNF2_N | 0.49 | 0.66 | 0.72 | 0.85 |
| AAA_33 | 0.29 | 0.50 | 0.72 | 0.69 |
| DEAD_4 | 0.23 | 0.26 | 0.72 | 0.31 |
| ResIII | 0.51 | 0.94 | 0.67 | 0.98 |
| NTPase_1 | 0.44 | 0.04 | 0.67 | 0.53 |
| NB-ARC_1st | 0.25 | 0.08 | 0.67 | 0.37 |
| Mur_ligase_M | 0.43 | 0.99 | 0.61 | 1.00 |
| Helicase_C_6 | 0.29 | 0.31 | 0.61 | 0.82 |
| AAA_19 | 0.23 | 0.74 | 0.61 | 0.78 |
| dNK_2 | 0.09 | 0.29 | 0.61 | 0.60 |
| AAA_12 | 0.50 | 0.34 | 0.61 | 0.58 |
| PEPCK_C | 0.11 | 0.25 | 0.61 | 0.54 |
| PEPCK_N | 0.11 | 0.25 | 0.61 | 0.53 |
| ATPase_2 | 0.16 | 0.05 | 0.61 | 0.42 |
| AAA_25 | 0.11 | 0.97 | 0.56 | 0.98 |
| MutS_V | 0.44 | 0.78 | 0.56 | 0.95 |
| Adenylsucc_synt | 0.46 | 0.87 | 0.56 | 0.93 |
| PEPCK_ATP_C | 0.26 | 0.45 | 0.56 | 0.74 |
| APS_kinase | 0.25 | 0.46 | 0.56 | 0.73 |
| PRK | 0.20 | 0.66 | 0.56 | 0.73 |
| APS_kinase_1 | 0.26 | 0.46 | 0.56 | 0.72 |
| Sulfotransfer_1 | 0.09 | 0.26 | 0.56 | 0.65 |
| TK_N | 0.14 | 0.49 | 0.56 | 0.48 |
| MG423 | 0.48 | 0.58 | 0.56 | 0.44 |
| SUA5 | 0.18 | 0.19 | 0.56 | 0.29 |
| Bac_DnaA_N | 0.12 | 0.96 | 0.50 | 0.98 |
| DnaB_C | 0.03 | 0.94 | 0.50 | 0.98 |
| UvrD_C_2 | 0.04 | 0.79 | 0.50 | 0.89 |
| UvrD-helicase | 0.24 | 0.70 | 0.50 | 0.79 |
| dNK | 0.05 | 0.27 | 0.50 | 0.58 |
| Terminase_6 | 0.09 | 0.10 | 0.50 | 0.53 |
| cobW | 0.32 | 0.51 | 0.50 | 0.45 |

| | | | | |
|---|---|---|---|---|
| AAA_11 | 0.22 | 0.11 | 0.50 | 0.44 |
| Helicase_RecD | 0.30 | 0.06 | 0.50 | 0.04 |
| DUF1726 | 0.28 | 0.04 | 0.50 | 0.02 |
| CoaE | 0.02 | 0.90 | 0.44 | 0.98 |
| Torsin | 0.06 | 0.95 | 0.44 | 0.97 |
| FtsK_SpoIIIE | 0.02 | 0.91 | 0.44 | 0.92 |
| FTHFS | 0.26 | 0.40 | 0.44 | 0.70 |
| AAA_26 | 0.36 | 0.56 | 0.44 | 0.68 |
| MobB | 0.32 | 0.25 | 0.44 | 0.60 |
| COG5410 | 0.03 | 0.04 | 0.44 | 0.38 |
| Guanylate_kin | 0.02 | 0.92 | 0.39 | 0.96 |
| DUF1846_N | 0.02 | 0.12 | 0.39 | 0.31 |
| PhoH | 0.06 | 0.86 | 0.33 | 0.91 |
| Sigma54_activ_2 | 0.00 | 0.41 | 0.33 | 0.82 |
| ArgK | 0.46 | 0.40 | 0.33 | 0.76 |
| RsgA_GTPase | 0.06 | 0.73 | 0.33 | 0.67 |
| PEPCK_ATP_N | 0.16 | 0.42 | 0.33 | 0.66 |
| AAA_32_2nd | 0.07 | 0.16 | 0.33 | 0.56 |
| SMC_N_2 | 0.02 | 0.15 | 0.33 | 0.56 |
| Helicase_C_8 | 0.06 | 0.21 | 0.33 | 0.53 |
| ABC_ATPase_C | 0.06 | 0.07 | 0.33 | 0.47 |
| DUF1611 | 0.28 | 0.10 | 0.33 | 0.33 |
| Terminase_3 | 0.01 | 0.08 | 0.33 | 0.26 |
| CPT | 0.01 | 0.06 | 0.33 | 0.24 |
| CLP1_P | 0.11 | 0.00 | 0.33 | 0.23 |
| Helicase_C_10 | 0.00 | 0.93 | 0.28 | 0.95 |
| SecA_DEAD | 0.00 | 0.92 | 0.28 | 0.95 |
| dNK_1 | 0.03 | 0.26 | 0.28 | 0.56 |
| CobU | 0.01 | 0.43 | 0.28 | 0.53 |
| PPK2 | 0.13 | 0.52 | 0.28 | 0.52 |
| Sulfotransfer_2 | 0.01 | 0.03 | 0.28 | 0.24 |
| Gtr1_RagA_N | 0.05 | 0.01 | 0.28 | 0.12 |
| AAA_28 | 0.01 | 0.17 | 0.22 | 0.24 |
| UvrD_C_2_1 | 0.00 | 0.06 | 0.22 | 0.18 |
| Thymidylate_kin_1 | 0.01 | 0.00 | 0.22 | 0.08 |
| GTP_HydF_2nd | 0.00 | 0.11 | 0.17 | 0.52 |

| | | | | |
|---|---|---|---|---|
| GTP_HydF_1st | 0.00 | 0.11 | 0.17 | 0.50 |
| KdpD | 0.03 | 0.32 | 0.17 | 0.40 |
| TrwB_AAD_bind | 0.05 | 0.27 | 0.17 | 0.38 |
| AAA_11_1 | 0.01 | 0.05 | 0.17 | 0.27 |
| NACHT | 0.00 | 0.02 | 0.17 | 0.19 |
| 6PF2K | 0.00 | 0.01 | 0.17 | 0.12 |
| TsaE | 0.00 | 0.92 | 0.11 | 0.96 |
| Cytidylate_kin_1 | 0.00 | 0.86 | 0.11 | 0.94 |
| Cytidylate_kin_2 | 0.00 | 0.86 | 0.11 | 0.94 |
| Cytidylate_kin_3 | 0.00 | 0.86 | 0.11 | 0.94 |
| Cytidylate_kin_4 | 0.00 | 0.86 | 0.11 | 0.94 |
| MlrC_C | 0.03 | 0.12 | 0.11 | 0.31 |
| CE15 | 0.00 | 0.06 | 0.11 | 0.24 |
| DUF927_C | 0.01 | 0.06 | 0.11 | 0.24 |
| Sulfotransfer_3 | 0.00 | 0.04 | 0.11 | 0.19 |
| AAA_6_N | 0.00 | 0.00 | 0.11 | 0.07 |
| Podovirus_Gp16 | 0.00 | 0.00 | 0.11 | 0.05 |
| AAA_7_N | 0.01 | 0.00 | 0.11 | 0.02 |
| Helicase_C_9 | 0.00 | 0.80 | 0.06 | 0.91 |
| LpxK_N | 0.00 | 0.49 | 0.06 | 0.74 |
| Hpr_kinase_C | 0.00 | 0.35 | 0.06 | 0.62 |
| Kinase-PPPase_C | 0.00 | 0.36 | 0.06 | 0.40 |
| Kinase-PPPase_N | 0.00 | 0.15 | 0.06 | 0.37 |
| Zeta_toxin | 0.00 | 0.03 | 0.06 | 0.16 |
| Dynamin_N | 0.00 | 0.00 | 0.06 | 0.09 |
| AAA_5 | 0.00 | 0.00 | 0.06 | 0.07 |
| P-mevalo_kinase | 0.00 | 0.01 | 0.06 | 0.06 |
| Viral_helicase1 | 0.00 | 0.00 | 0.06 | 0.05 |
| AAA_8 | 0.00 | 0.00 | 0.06 | 0.03 |
| P-mevalo_kinase_1 | 0.00 | 0.00 | 0.06 | 0.03 |
| AAA_9_N | 0.00 | 0.00 | 0.06 | 0.02 |
| Dynein_heavy_1st | 0.00 | 0.00 | 0.06 | 0.02 |
| AAA_22_like | 0.00 | 0.00 | 0.06 | 0.01 |
| DUF257 | 0.03 | 0.00 | 0.06 | 0.00 |
| TIP49_1st | 0.00 | 0.00 | 0.06 | 0.00 |

| | | | | |
|---|---|---|---|---|
| DNA_pol3_delta_N | 0.00 | 0.62 | 0.00 | 0.76 |
| UvrD-helicase_1 | 0.00 | 0.17 | 0.00 | 0.30 |
| Exonuc_V_gamma | 0.00 | 0.19 | 0.00 | 0.25 |
| Helicase_C_7 | 0.00 | 0.10 | 0.00 | 0.21 |
| Sulphotransf | 0.00 | 0.03 | 0.00 | 0.15 |
| Zot | 0.00 | 0.02 | 0.00 | 0.15 |
| DNA_pol3_chi | 0.00 | 0.27 | 0.00 | 0.11 |
| IPT | 0.00 | 0.01 | 0.00 | 0.09 |
| Cas_Csn2 | 0.00 | 0.02 | 0.00 | 0.08 |
| MukB | 0.00 | 0.03 | 0.00 | 0.08 |
| HDA2-3 | 0.00 | 0.00 | 0.00 | 0.04 |
| Septin | 0.00 | 0.00 | 0.00 | 0.04 |
| Cas_St_Csn2 | 0.00 | 0.00 | 0.00 | 0.03 |
| Kinesin | 0.00 | 0.00 | 0.00 | 0.02 |
| Myosin_head | 0.00 | 0.00 | 0.00 | 0.02 |
| Myosin_head_1 | 0.00 | 0.00 | 0.00 | 0.02 |
| Myosin_head_2 | 0.00 | 0.00 | 0.00 | 0.02 |
| Parvo_NS1 | 0.00 | 0.00 | 0.00 | 0.02 |
| PRK10689 | 0.00 | 0.10 | 0.00 | 0.02 |
| PRK13709 | 0.00 | 0.00 | 0.00 | 0.02 |
| SulA | 0.00 | 0.08 | 0.00 | 0.02 |
| DENN | 0.00 | 0.00 | 0.00 | 0.01 |
| Herpes_TK | 0.00 | 0.00 | 0.00 | 0.01 |
| PRK07993_1st | 0.00 | 0.01 | 0.00 | 0.01 |
| RHD3_1st | 0.00 | 0.00 | 0.00 | 0.01 |
| AAA_33_like | 0.00 | 0.00 | 0.00 | 0.00 |
| AAA_33,ADK_lid | 0.00 | 0.00 | 0.00 | 0.00 |
| AAA_33,RNA_pol_Rpc34 | 0.00 | 0.00 | 0.00 | 0.00 |
| ABC_tran_3 | 0.00 | 0.00 | 0.00 | 0.00 |
| Adaptin_binding | 0.00 | 0.00 | 0.00 | 0.00 |
| ADK_lid,Cytidylate_kin_2 | 0.00 | 0.00 | 0.00 | 0.00 |
| ATP-synt_ab,ATP-synt_ab_C | 0.00 | 0.00 | 0.00 | 0.00 |
| ATP-synt_ab,ATP-synt_ab_N | 0.00 | 0.00 | 0.00 | 0.00 |
| CDC73_C | 0.00 | 0.00 | 0.00 | 0.00 |

| | | | | |
|---|---|---|---|---|
| CENP-M | 0.00 | 0.00 | 0.00 | 0.00 |
| COG5177 | 0.00 | 0.00 | 0.00 | 0.00 |
| CSM2 | 0.00 | 0.00 | 0.00 | 0.00 |
| Cytidylate_kin_2,Thymidylate_kin | 0.00 | 0.00 | 0.00 | 0.00 |
| DAP3_C,DAP3_N | 0.00 | 0.00 | 0.00 | 0.00 |
| DAP3_N | 0.00 | 0.00 | 0.00 | 0.00 |
| DEAD_1,KOG0951_2nd | 0.00 | 0.00 | 0.00 | 0.00 |
| DLIC | 0.00 | 0.00 | 0.00 | 0.00 |
| Elong_Iki1 | 0.00 | 0.00 | 0.00 | 0.00 |
| ELP6 | 0.00 | 0.00 | 0.00 | 0.00 |
| EUF08601 | 0.00 | 0.00 | 0.00 | 0.00 |
| F_UNCLASSIFIED | 0.00 | 0.00 | 0.00 | 0.00 |
| Flavi_DEAD | 0.00 | 0.00 | 0.00 | 0.00 |
| Folliculin_C | 0.00 | 0.00 | 0.00 | 0.00 |
| GBP | 0.00 | 0.00 | 0.00 | 0.00 |
| GBP_1 | 0.00 | 0.00 | 0.00 | 0.00 |
| HA2_N,DEAD_3 | 0.00 | 0.00 | 0.00 | 0.00 |
| IPPT,IPT | 0.00 | 0.00 | 0.00 | 0.00 |
| KOG0163_1,Myosin_head_1 | 0.00 | 0.00 | 0.00 | 0.00 |
| KOG0384_1,Helicase_C | 0.00 | 0.00 | 0.00 | 0.00 |
| KOG2538_1st | 0.00 | 0.00 | 0.00 | 0.00 |
| LAP1C | 0.00 | 0.00 | 0.00 | 0.00 |
| Microtub_bd | 0.00 | 0.00 | 0.00 | 0.00 |
| MobB_1 | 0.00 | 0.00 | 0.00 | 0.00 |
| Myosin_head,IQ_2 | 0.00 | 0.00 | 0.00 | 0.00 |
| NTPase_P4 | 0.00 | 0.00 | 0.00 | 0.00 |
| Orbi_VP4_1st | 0.00 | 0.00 | 0.00 | 0.00 |
| ORC2_1st | 0.00 | 0.00 | 0.00 | 0.00 |
| ORC3_N_1st | 0.00 | 0.00 | 0.00 | 0.00 |
| PAXNEB | 0.00 | 0.00 | 0.00 | 0.00 |
| Polyoma_lg_T_C | 0.00 | 0.00 | 0.00 | 0.00 |
| PRK11000,ABC_tran | 0.00 | 0.00 | 0.00 | 0.00 |
| PRK11000,ABC_tran_1 | 0.00 | 0.00 | 0.00 | 0.00 |
| PSY3 | 0.00 | 0.00 | 0.00 | 0.00 |
| Rap_GAP | 0.00 | 0.00 | 0.00 | 0.00 |
| Sigma54_activat,CDC48 | 0.00 | 0.00 | 0.00 | 0.00 |

| | | | | |
|---|---|---|---|---|
| Sigma54_activat,KOG0729 | 0.00 | 0.00 | 0.00 | 0.00 |
| Sigma54_activat,Vps4_C_C | 0.00 | 0.00 | 0.00 | 0.00 |
| Torsin,RecA | 0.00 | 0.00 | 0.00 | 0.00 |
| UNK_F_TYPE | 0.00 | 0.00 | 0.00 | 0.00 |
| UvrA,ABC_tran | 0.00 | 0.00 | 0.00 | 0.00 |

[a] Distributions were calculated as described in the main text. The seven representative families chosen for further analysis are written in red, with the selection criteria provided in the main text.

**Table S2**. Representative archaeal and bacterial NTPases studied in this work.[a]

| Reference Domain | F-Group Name | Organism | %Identity (%Positives) | Strand Order | Walker A Sequence | Res. (Å) | Ligand |
|---|---|---|---|---|---|---|---|
| e2it1A8 | ABC_tran | *Pyrococcus horikoshii* | 47 (69) | 23415(6) | GPSGSGKS | 1.94 | None (ATP)[b] |
| e3ievA2 | MnmE_helical_2nd | *Aquifex aeolicus* | 44 (61) | (2)31456 | GKPNVGKS | 1.9 | GNP |
| e3k1jA2 | Sigma54_activat | *Thermococcus onnurineus* | 40 (54) | 23415 | GEPGTGKS | 2 | ADP |
| e6ojxA1 | T2SSe | *Geobacter metallireducens* | 40 (56) | 324516(7) | GPTGSGKS | 1.89 | ATP |
| e5jrjA2 | RecA | *Herbaspirillum seropedicae* | 71 (85) | 324516(7)8 | GPESSGKT | 1.7 | ATP /ADP |
| e2qy9A2 | SRP54 | *Escherichia coli* | 57 (72) | 3241567 | GVNGVGKT | 1.9 | None (GTP)[c] |

[a] For details of how these NTPases were selected, see the main and **Table S1**. %Identity and %Positives are relative to consensus hmmemit. For the strand order, ()=antiparallel. [b] While there is no ligand in the associated structure, Uniprot (11) lists the P-loop NTPase PH0203 from *Pyrococcus horikoshii* as a 362aa-long hypothetical maltose/maltodextrin ATP-binding protein. [c] While there is no ligand in the associated structure, the *E. coli* SRP-receptor FtsY (PDB ID: 2QY9(12)) has GTPase activity (13).

**Table S3**. Selected control sequences studied in this work

| Reference Domain | X-Group Name | Organism | Control Seqeunce | Res. (Å) | Ligand |
|---|---|---|---|---|---|
| e1mt5A1 | X-Group 7552 | *Rattus norvegicus* | KSPGGSSG | 2.30 | None |
| e5dmyA4 | Immunoglobulin-like beta-sandwich | *Bifidobacterium bifidum* | KGSGNKVP | 1.95 | None |
| e4bkrA1 | Rossmann-like | *Yersinia pestis A1122* | EGKPGTSG | 1.80 | None |
| e1re5D1 | L-Aspartase middle domain-like | *Pseudomonas putida KT2440* | PGKGGSST | 2.60 | None |
| e4lnbA1 | Repetitive alpha hairpins | *Aspergillus fumigatus Af293* | TGPSSEKG | 1.75 | None |
| e3o26A1 | Rossmann-like | *Papaver somniferum* | VVTGGNKG | 1.91 | NDP |

**Table S4.** The ff99SB-ILDN (14) force field parameters used to describe the ligand $HPO_4^{2-}$.

| **Atom Name** | **Atom Type** | **Partial Charge** | **Atom Name** | **Atom Type** | **Partial Charge** |
|---|---|---|---|---|---|
| P | PP | 1.320903 | O3 | O2P | -0.956501 |
| O1 | OHP | -0.815401 | O4 | O2P | -0.956501 |
| O2 | O2P | -0.956501 | HOP | HOP | 0.364000 |

**Table S5.** The ff99SB-ILDN (14) force field parameters used to describe the ligand GTP.

| Atom Name | Atom Type | Partial Charge | Atom Name | Atom Type | Partial Charge |
|---|---|---|---|---|---|
| O1G | O3 | -0.9526 | C8 | CK | 0.1374 |
| PG | P | 1.2650 | H8 | H5 | 0.1640 |
| O2G | O3 | -0.9526 | N7 | NB | -0.5709 |
| O3G | O3 | -0.9526 | C5 | CB | 0.1744 |
| O3B | OS | -0.5322 | C6 | C | 0.4770 |
| PB | P | 1.3852 | O6 | O | -0.5597 |
| O1B | O2 | -0.8894 | N1 | NA | -0.4787 |
| O2B | O2 | -0.8894 | H1 | H | 0.3424 |
| O3A | OS | -0.5689 | C2 | CA | 0.7657 |
| PA | P | 1.2532 | N2 | N2 | -0.9672 |
| O1A | O2 | -0.8799 | H21 | H | 0.4364 |
| O2A | O2 | -0.8799 | H22 | H | 0.4364 |
| O5' | OS | -0.5987 | N3 | NC | -0.6323 |
| C5' | CT | 0.0558 | C4 | CB | 0.1222 |
| H5'1 | H1 | 0.0679 | C3' | CT | 0.2022 |
| H5'2 | H1 | 0.0679 | O3' | OH | -0.6541 |
| C4' | CT | 0.1065 | H3T | HO | 0.4376 |
| H4' | H1 | 0.1174 | H3' | H1 | 0.0615 |
| O4' | OS | -0.3548 | C2' | CT | 0.0670 |
| C1' | CT | 0.0191 | H2'1 | H1 | 0.0972 |
| H1' | H2 | 0.2006 | O2' | OH | -0.6139 |
| N9 | N* | 0.0492 | HO'2 | HO | 0.4186 |

**Table S6**. Residues that interact with the $Mg^{2+}$ ion in the representative P-Loop structures.

| **Residue** | G | X | X | G | X | G | K | S/T |
|---|---|---|---|---|---|---|---|---|
| **Count** | 1 | 0 | 5 | 2 | 1 | 1 | 4 | 419 |

**Table S7**. Occurrence (% simulation time) of correlated stretches of $\alpha_L$ and $\alpha_R$ conformations in Walker-A - derived octapeptides

| Sequence | 4-5 (RL) | 5-6 (RL) | 6-7 (RL) | 4-5 (LR) | 5-6 (LR) | 6-7 (LR) | 4-6 (RLR) | 5-7 (RLR) | 4-6 (LRL) | 5-7 (LRL) | 4-7 (LRLR) |
|---|---|---|---|---|---|---|---|---|---|---|---|
| **Unliganded** | | | | | | | | | | | |
| **SGAGKT** | 0.22 | 2.63 | 0.55 | 1.03 | 0.13 | 5.33 | 0.02 | 0.72 | 0.09 | 0.02 | 0.02 |
| **GPSGSGKS** | 0.80 | 7.76 | 0.61 | 4.57 | 0.65 | 6.77 | 0.01 | 1.68 | 1.23 | 0.00 | 0.22 |
| **GKPNVGKS** | 1.72 | 5.41 | 0.45 | 7.07 | 0.06 | 3.74 | 0.01 | 0.63 | 1.72 | 0.00 | 0.25 |
| **GEPGTGKS** | 0.23 | 17.59 | 0.63 | 16.60 | 0.18 | 7.49 | 0.04 | 2.80 | 7.10 | 0.06 | 1.16 |
| **GPTGSGKS** | 0.81 | 8.66 | 0.68 | 4.57 | 0.68 | 7.20 | 0.02 | 1.83 | 1.15 | 0.09 | 0.23 |
| **GPESSGKT** | 9.91 | 11.08 | 0.96 | 1.30 | 0.46 | 11.57 | 0.13 | 2.25 | 0.48 | 0.17 | 0.19 |
| **GVNGVGKT** | 0.39 | 6.69 | 0.47 | 9.58 | 0.05 | 6.04 | 0.00 | 1.24 | 1.50 | 0.03 | 0.30 |
| **$HPO_4^{2-}$** | | | | | | | | | | | |
| **SGAGKT** | 0.40 | 2.48 | 0.50 | 1.72 | 0.19 | 6.32 | 0.00 | 0.70 | 0.19 | 0.03 | 0.04 |
| **GPSGSGKS** | 0.94 | 7.03 | 0.67 | 4.79 | 0.64 | 7.14 | 0.06 | 1.76 | 1.14 | 0.10 | 0.22 |
| **GKPNVGKS** | 1.51 | 5.33 | 0.45 | 6.03 | 0.13 | 5.07 | 0.07 | 1.21 | 1.64 | 0.00 | 0.60 |
| **GEPGTGKS** | 0.26 | 15.55 | 0.70 | 16.11 | 0.18 | 7.41 | 0.01 | 2.92 | 6.85 | 0.00 | 1.27 |
| **GPTGSGKS** | 0.56 | 7.42 | 0.71 | 4.80 | 0.80 | 7.77 | 0.00 | 1.79 | 1.09 | 0.04 | 0.23 |
| **GPESSGKT** | 8.15 | 14.43 | 0.92 | 3.66 | 0.54 | 13.08 | 0.03 | 3.33 | 1.52 | 0.08 | 0.40 |
| **GVNGVGKT** | 0.59 | 6.00 | 0.95 | 10.17 | 0.05 | 6.28 | 0.00 | 1.28 | 1.79 | 0.00 | 0.49 |
| **GTP** | | | | | | | | | | | |
| **SGAGKT** | 0.49 | 3.54 | 0.78 | 5.53 | 0.41 | 13.57 | 0.02 | 2.28 | 1.32 | 0.00 | 0.98 |
| **GPSGSGKS** | 0.49 | 3.54 | 0.78 | 5.53 | 0.41 | 13.57 | 0.02 | 2.28 | 1.32 | 0.00 | 0.98 |
| **GKPNVGKS** | 0.41 | 6.12 | 0.31 | 20.49 | 1.05 | 7.46 | 0.24 | 3.49 | 1.72 | 0.14 | 0.91 |
| **GEPGTGKS** | 2.36 | 3.28 | 0.53 | 11.79 | 0.24 | 4.06 | 0.00 | 0.66 | 0.64 | 0.00 | 0.29 |
| **GPTGSGKS** | 0.34 | 1.51 | 0.06 | 33.53 | 2.19 | 0.96 | 0.34 | 0.32 | 0.98 | 0.00 | 0.21 |
| **GPESSGKT** | 0.22 | 4.26 | 0.15 | 26.42 | 0.13 | 4.37 | 0.00 | 0.91 | 1.02 | 0.01 | 0.09 |
| **GVNGVGKT** | 6.16 | 19.97 | 0.33 | 3.18 | 1.89 | 6.02 | 1.62 | 1.42 | 0.64 | 0.04 | 0.09 |

**Table S8**. Peptide-ligand hydrogen bonding (% simulation time) between the Walker-A P-loop sequences studied in this work. in complex with either $HPO_4^{2-}$ or GTP.[a]

| Sequence | | $HPO_4^{2-}$ SGAGKT | $HPO_4^{2-}$ GPSGSGKS | $HPO_4^{2-}$ GKPNVGKS | $HPO_4^{2-}$ GEPGTGKS | $HPO_4^{2-}$ GPTGSGKS | $HPO_4^{2-}$ GPESSGKT | $HPO_4^{2-}$ GVNGVGKT | GTP-α SGAGKT | GTP-α GPSGSGKS | GTP-α GKPNVGKS | GTP-α GEPGTGKS | GTP-α GPTGSGKS | GTP-α GPESSGKT | GTP-α GVNGVGKT |
|---|---|---|---|---|---|---|---|---|---|---|---|---|---|---|---|
| | **N** | 12.81 | 12.78 | 12.95 | 9.36 | 11.54 | 10.51 | 12.33 | 35.50 | 46.21 | 41.21 | 35.03 | 40.08 | 29.71 | 42.76 |
| **1** | **R** | | 0.00 | 0.00 | 0.00 | 0.00 | 0.00 | 0.00 | | 0.00 | 0.00 | 0.00 | 0.00 | 0.00 | 0.00 |
| | **O** | | 0.12 | 0.13 | 0.08 | 0.09 | 0.05 | 0.09 | | 0.00 | 0.00 | 0.00 | 0.00 | 0.00 | 0.00 |
| | **N** | | 0.00 | 3.54 | 0.97 | 0.00 | 0.00 | 3.19 | | 0.00 | 19.61 | 0.62 | 0.00 | 0.00 | 14.86 |
| **2** | **R** | | 0.00 | 17.15 | 0.02 | 0.00 | 0.00 | 0.00 | | 0.00 | 25.05 | 0.00 | 0.00 | 0.00 | 0.00 |
| | **O** | | 0.03 | 0.17 | 0.09 | 0.03 | 0.01 | 0.06 | | 0.00 | 0.00 | 0.00 | 0.00 | 0.00 | 0.00 |
| | **N** | | 1.52 | 0.00 | 0.00 | 0.82 | 0.19 | 3.50 | | 3.27 | 0.00 | 0.00 | 0.38 | 0.04 | 16.81 |
| **3** | **R** | 7.91 | 3.35 | 0.00 | 0.00 | 1.56 | 0.02 | 4.40 | 8.17 | 4.72 | 0.00 | 0.00 | 0.52 | 0.00 | 13.71 |
| | **O** | 0.10 | 0.02 | 0.00 | 0.00 | 0.01 | 0.00 | 0.08 | 0.00 | 0.00 | 0.00 | 0.00 | 0.00 | 0.00 | 0.00 |
| | **N** | 7.54 | 0.78 | 0.19 | 0.15 | 0.55 | 0.22 | 1.66 | 19.71 | 2.07 | 0.01 | 0.00 | 0.21 | 0.02 | 10.36 |
| **4** | **R** | 0.00 | 0.00 | 0.96 | 0.00 | 0.00 | 5.20 | 0.00 | 0.00 | 0.00 | 1.53 | 0.00 | 0.00 | 12.15 | 0.00 |
| | **O** | 0.09 | 0.01 | 0.00 | 0.00 | 0.02 | 0.01 | 0.03 | 0.00 | 0.00 | 0.00 | 0.00 | 0.00 | 0.00 | 0.00 |
| | **N** | 1.28 | 1.32 | 0.44 | 0.52 | 1.58 | 0.51 | 0.33 | 4.47 | 4.59 | 0.08 | 0.00 | 3.06 | 0.08 | 1.87 |
| **5** | **R** | 0.00 | 4.54 | 0.00 | 0.74 | 4.36 | 4.41 | 0.00 | 0.00 | 7.84 | 0.00 | 0.12 | 8.36 | 17.45 | 0.00 |
| | **O** | 0.03 | 0.01 | 0.01 | 0.01 | 0.02 | 0.01 | 0.01 | 0.00 | 0.00 | 0.00 | 0.00 | 0.00 | 0.00 | 0.00 |
| | **N** | 0.61 | 0.80 | 0.36 | 0.22 | 1.06 | 0.36 | 0.36 | 4.33 | 8.38 | 0.00 | 0.32 | 5.60 | 0.55 | 2.36 |
| **6** | **R** | 0.00 | 0.00 | 0.00 | 0.00 | 0.00 | 0.00 | 0.00 | 0.00 | 0.00 | 0.00 | 0.00 | 0.00 | 0.00 | 0.00 |
| | **O** | 0.03 | 0.03 | 0.01 | 0.01 | 0.03 | 0.02 | 0.01 | 0.00 | 0.00 | 0.00 | 0.00 | 0.00 | 0.00 | 0.00 |
| | **N** | 0.61 | 0.72 | 0.50 | 0.36 | 0.70 | 0.37 | 0.52 | 0.28 | 9.85 | 4.23 | 0.43 | 2.75 | 1.74 | 2.55 |
| **7** | **R** | 17.88 | 17.32 | 11.49 | 14.46 | 15.22 | 14.13 | 14.01 | 37.28 | 44.46 | 33.81 | 59.42 | 48.66 | 47.39 | 42.45 |
| | **O** | 0.02 | 0.02 | 0.01 | 0.01 | 0.02 | 0.01 | 0.01 | 0.00 | 0.00 | 0.00 | 0.00 | 0.00 | 0.00 | 0.00 |
| | **N** | 0.98 | 1.19 | 0.69 | 0.53 | 0.99 | 0.32 | 0.53 | 0.46 | 5.82 | 2.99 | 0.59 | 1.04 | 1.74 | 0.64 |
| **8** | **R** | 1.33 | 3.73 | 1.98 | 1.99 | 2.91 | 0.48 | 0.90 | 0.67 | 8.40 | 4.07 | 0.86 | 1.62 | 2.17 | 1.32 |
| | **O** | 0.01 | 0.01 | 0.01 | 0.01 | 0.01 | 0.01 | 0.01 | 0.00 | 0.00 | 0.00 | 0.00 | 0.00 | 0.00 | 0.00 |

| Sequence | | GTP-β SGAGKT | GTP-β GPSGSGKS | GTP-β GKPNVGKS | GTP-β GEPGTGKS | GTP-β GPTGSGKS | GTP-β GPESSGKT | GTP-β GVNGVGKT | GTP-γ SGAGKT | GTP-γ GPSGSGKS | GTP-γ GKPNVGKS | GTP-γ GEPGTGKS | GTP-γ GPTGSGKS | GTP-γ GPESSGKT | GTP-γ GVNGVGKT |
|---|---|---|---|---|---|---|---|---|---|---|---|---|---|---|---|
| | N | 22.72 | 31.42 | 26.07 | 46.45 | 39.77 | 26.57 | 30.86 | 36.46 | 43.07 | 41.97 | 44.31 | 46.37 | 43.95 | 53.56 |
| 1 | R | | 0.00 | 0.00 | 0.00 | 0.00 | 0.00 | 0.00 | | 0.00 | 0.00 | 0.00 | 0.00 | 0.00 | 0.00 |
| | O | | 0.00 | 0.00 | 0.00 | 0.00 | 0.00 | 0.00 | | 0.00 | 0.00 | 0.00 | 0.00 | 0.00 | 0.00 |
| | N | | 0.00 | 3.31 | 0.03 | 0.00 | 0.00 | 1.57 | | 0.00 | 6.46 | 0.16 | 0.00 | 0.00 | 8.24 |
| 2 | R | | 0.00 | 11.86 | 0.00 | 0.00 | 0.00 | 0.00 | | 0.00 | 30.11 | 0.00 | 0.00 | 0.00 | 0.00 |
| | O | | 0.00 | 0.00 | 0.00 | 0.00 | 0.00 | 0.00 | | 0.00 | 0.00 | 0.00 | 0.00 | 0.00 | 0.00 |
| | N | | 0.20 | 0.00 | 0.00 | 0.16 | 0.10 | 1.19 | | 1.37 | 0.00 | 0.00 | 1.56 | 0.00 | 8.40 |
| 3 | R | 4.17 | 0.28 | 0.00 | 0.00 | 0.14 | 0.00 | 1.63 | 25.24 | 2.32 | 0.00 | 0.00 | 1.49 | 0.00 | 8.47 |
| | O | 0.00 | 0.00 | 0.00 | 0.00 | 0.00 | 0.00 | 0.00 | 0.00 | 0.00 | 0.00 | 0.00 | 0.00 | 0.00 | 0.00 |
| | N | 6.24 | 0.18 | 0.01 | 0.01 | 0.20 | 0.12 | 0.78 | 16.28 | 0.09 | 0.00 | 0.00 | 1.06 | 0.01 | 3.00 |
| 4 | R | 0.00 | 0.00 | 0.19 | 0.00 | 0.00 | 3.79 | 0.00 | 0.00 | 0.00 | 0.10 | 0.00 | 0.00 | 29.74 | 0.00 |
| | O | 0.00 | 0.00 | 0.00 | 0.00 | 0.00 | 0.00 | 0.00 | 0.00 | 0.00 | 0.00 | 0.00 | 0.00 | 0.00 | 0.00 |
| | N | 4.62 | 1.68 | 0.00 | 0.00 | 0.25 | 0.03 | 0.01 | 2.04 | 0.25 | 0.00 | 0.00 | 1.19 | 0.03 | 0.00 |
| 5 | R | 0.00 | 1.93 | 0.00 | 0.01 | 1.60 | 14.77 | 0.00 | 0.00 | 3.24 | 0.00 | 0.01 | 2.62 | 5.57 | 0.00 |
| | O | 0.00 | 0.00 | 0.00 | 0.00 | 0.00 | 0.00 | 0.00 | 0.00 | 0.00 | 0.00 | 0.00 | 0.00 | 0.00 | 0.00 |
| | N | 3.31 | 0.30 | 0.00 | 0.00 | 0.93 | 0.54 | 0.06 | 1.55 | 0.03 | 0.00 | 0.00 | 0.22 | 0.19 | 0.00 |
| 6 | R | 0.00 | 0.00 | 0.00 | 0.00 | 0.00 | 0.00 | 0.00 | 0.00 | 0.00 | 0.00 | 0.00 | 0.00 | 0.00 | 0.00 |
| | O | 0.00 | 0.00 | 0.00 | 0.00 | 0.00 | 0.00 | 0.00 | 0.00 | 0.00 | 0.00 | 0.00 | 0.00 | 0.00 | 0.00 |
| | N | 1.22 | 0.48 | 0.97 | 0.00 | 0.25 | 0.37 | 0.75 | 1.42 | 4.23 | 0.03 | 0.00 | 0.03 | 0.23 | 10.32 |
| 7 | R | 25.08 | 28.70 | 20.85 | 16.33 | 20.28 | 30.89 | 33.06 | 48.39 | 66.07 | 42.00 | 82.98 | 68.96 | 35.38 | 62.11 |
| | O | 0.00 | 0.00 | 0.00 | 0.00 | 0.00 | 0.00 | 0.00 | 0.00 | 0.00 | 0.00 | 0.00 | 0.00 | 0.00 | 0.00 |
| | N | 0.22 | 1.65 | 0.12 | 0.08 | 0.06 | 0.19 | 0.01 | 0.16 | 3.94 | 0.55 | 31.01 | 10.76 | 0.29 | 9.69 |
| 8 | R | 0.51 | 2.87 | 0.90 | 0.35 | 0.22 | 0.19 | 1.03 | 0.22 | 5.76 | 0.98 | 34.79 | 12.71 | 0.69 | 10.08 |
| | O | 0.00 | 0.00 | 0.00 | 0.00 | 0.00 | 0.00 | 0.00 | 0.00 | 0.00 | 0.00 | 0.00 | 0.00 | 0.00 | 0.00 |

| Sequence | | SGAGKT | GPSGSGKS | GKPNVGKS | GEPGTGKS | GPTGSGKS | GPESSGKT | GVNGVGKT | SGAGKT | GPSGSGKS | GKPNVGKS | GEPGTGKS | GPTGSGKS | GPESSGKT | GVNGVGKT |
|---|---|---|---|---|---|---|---|---|---|---|---|---|---|---|---|
| | | GTP-triphosphate | | | | | | | GTP base+sugar | | | | | | |
| 1 | N | 68.86 | 75.47 | 69.85 | 80.01 | 77.66 | 68.97 | 83.31 | 27.19 | 51.15 | 44.83 | 77.26 | 65.40 | 23.86 | 49.28 |
| | R | | 0.00 | 0.00 | 0.00 | 0.00 | 0.00 | 0.00 | | 0.00 | 0.00 | 0.00 | 0.00 | 0.00 | 0.00 |
| | O | | 0.00 | 0.00 | 0.00 | 0.00 | 0.00 | 0.00 | | 1.68 | 2.48 | 0.31 | 1.60 | 0.49 | 1.06 |
| 2 | N | | 0.00 | 28.91 | 0.81 | 0.00 | 0.00 | 24.58 | | 0.00 | 4.25 | 0.73 | 0.00 | 0.00 | 1.66 |
| | R | | 0.00 | 42.90 | 0.00 | 0.00 | 0.00 | 0.00 | | 0.00 | 3.81 | 0.61 | 0.00 | 0.00 | 0.00 |
| | O | | 0.00 | 0.00 | 0.00 | 0.00 | 0.00 | 0.00 | | 6.00 | 13.10 | 3.61 | 2.13 | 0.53 | 1.81 |
| 3 | N | | 4.82 | 0.00 | 0.00 | 2.06 | 0.14 | 26.35 | | 3.08 | 0.00 | 0.00 | 0.84 | 0.43 | 1.04 |
| | R | 37.05 | 7.30 | 0.00 | 0.00 | 2.14 | 0.00 | 23.48 | 1.22 | 1.18 | 0.00 | 0.00 | 1.25 | 3.14 | 3.06 |
| | O | 0.00 | 0.00 | 0.00 | 0.00 | 0.00 | 0.00 | 0.00 | 1.69 | 0.89 | 1.43 | 0.09 | 1.07 | 0.58 | 31.99 |
| 4 | N | 40.16 | 2.32 | 0.02 | 0.01 | 1.43 | 0.16 | 13.95 | 8.84 | 8.27 | 15.85 | 26.86 | 18.61 | 0.71 | 2.90 |
| | R | 0.00 | 0.00 | 1.77 | 0.00 | 0.00 | 45.34 | 0.00 | 0.00 | 0.00 | 3.41 | 0.00 | 0.00 | 1.58 | 0.00 |
| | O | 0.00 | 0.00 | 0.00 | 0.00 | 0.00 | 0.00 | 0.00 | 3.27 | 3.66 | 0.62 | 0.11 | 1.89 | 1.15 | 19.37 |
| 5 | N | 10.77 | 4.80 | 0.08 | 0.01 | 4.33 | 0.14 | 1.88 | 2.80 | 6.70 | 16.39 | 33.67 | 20.00 | 0.79 | 2.16 |
| | R | 0.00 | 12.80 | 0.00 | 0.14 | 12.49 | 37.49 | 0.00 | 0.00 | 4.53 | 0.00 | 5.34 | 3.54 | 2.00 | 0.00 |
| | O | 0.00 | 0.00 | 0.00 | 0.00 | 0.00 | 0.00 | 0.00 | 3.81 | 4.03 | 2.45 | 50.60 | 19.05 | 5.66 | 2.44 |
| 6 | N | 7.87 | 8.66 | 0.01 | 0.32 | 6.19 | 1.05 | 2.39 | 3.21 | 15.47 | 13.29 | 24.24 | 25.20 | 2.32 | 4.40 |
| | R | 0.00 | 0.00 | 0.00 | 0.00 | 0.00 | 0.00 | 0.00 | 0.00 | 0.00 | 0.00 | 0.00 | 0.00 | 0.00 | 0.00 |
| | O | 0.00 | 0.00 | 0.00 | 0.00 | 0.00 | 0.00 | 0.00 | 3.85 | 13.46 | 2.85 | 66.02 | 34.92 | 1.46 | 3.46 |
| 7 | N | 2.86 | 14.54 | 5.10 | 0.43 | 3.03 | 2.32 | 13.48 | 5.34 | 2.93 | 3.67 | 1.65 | 4.30 | 2.87 | 15.29 |
| | R | 68.39 | 80.03 | 56.55 | 90.81 | 82.19 | 71.95 | 77.50 | 12.56 | 14.74 | 6.74 | 37.85 | 25.25 | 27.17 | 5.47 |
| | O | 0.00 | 0.00 | 0.00 | 0.00 | 0.00 | 0.00 | 0.00 | 2.63 | 1.86 | 1.92 | 0.33 | 1.01 | 0.70 | 1.44 |
| 8 | N | 0.83 | 11.31 | 3.64 | 31.67 | 11.85 | 2.09 | 10.35 | 5.11 | 2.08 | 1.83 | 0.28 | 1.49 | 1.70 | 3.94 |
| | R | 1.38 | 16.44 | 5.91 | 35.79 | 14.44 | 3.01 | 11.93 | 5.61 | 3.87 | 3.17 | 0.80 | 2.19 | 2.41 | 4.85 |
| | O | 0.00 | 0.00 | 0.00 | 0.00 | 0.00 | 0.00 | 0.00 | 15.85 | 11.21 | 13.30 | 3.17 | 7.79 | 34.90 | 21.03 |

[a]Analysis based on snapshots taken every 10 ps of the simulation, calculated as described in the main text. The residue numbering of the SGAGKT hexapeptide has been shifted by +2 to align with the other peptides presented here for clarity. This data is shown visually in **Figures 2** and **3**.

**Table S9**. Peptide-ligand hydrogen bonding (% simulation time) between the peptides studied in this work. in complex with either $HPO_4^{2-}$ or GTP.[a]

| | $HPO_4^{2-}$ | GTP $P_\alpha$ | GTP $P_\beta$ | GTP $P_\gamma$ | GTP triphosphate | GTP sugar+base | GTP whole |
|---|---|---|---|---|---|---|---|
| **SGAGKT** | 36.50 | 76.50 | 48.98 | 79.23 | 97.30 | 70.21 | 99.20 |
| **GPSGSGKS** | 34.99 | 78.94 | 54.40 | 84.61 | 97.60 | 80.62 | 99.46 |
| **GKPNVGKS** | 39.91 | 79.48 | 51.36 | 83.48 | 97.82 | 77.28 | 99.56 |
| **GEPGTGKS** | 24.73 | 79.51 | 57.91 | 93.56 | 98.89 | 97.89 | 99.90 |
| **GPTGSGKS** | 30.83 | 77.34 | 55.00 | 87.20 | 98.03 | 87.20 | 99.61 |
| **GPESSGKT** | 27.63 | 75.37 | 57.09 | 79.60 | 96.60 | 74.36 | 98.77 |
| **GVNGVGKT** | 28.71 | 81.98 | 58.66 | 86.23 | 98.15 | 85.15 | 99.66 |
| **KSPGGSSG** | 49.14 | 84.15 | 43.89 | 88.54 | 99.29 | 77.57 | 99.81 |
| **KGSGNKVP** | 53.04 | 86.67 | 58.35 | 87.06 | 99.26 | 84.26 | 99.87 |
| **EGKPGTSG** | 25.17 | 73.71 | 43.70 | 72.72 | 94.69 | 76.11 | 98.32 |
| **PGKGGSST** | 38.57 | 79.26 | 52.08 | 78.30 | 97.28 | 66.89 | 99.18 |
| **TGPSSEKG** | 32.25 | 76.88 | 50.52 | 83.45 | 98.29 | 89.79 | 99.71 |
| **VVTGGNKG** | 25.48 | 80.85 | 47.55 | 72.05 | 95.98 | 75.51 | 98.89 |

[a]Analysis based on snapshots taken every 10 ps of the simulation. calculated as described in the main text. Sequences from GTP-binding NTPases are shown in green and sequences from ATP-binding NTPases are shown in blue.

**Table S10**. Occurrence (% simulation time) of correlated stretches of $\alpha_L$ and $\alpha_R$ conformations in Walker A-derived octapeptides during simulations performed using the CHARMM36m force field.(10)

| Sequence | 4-5 (RL) | 5-6 (RL) | 6-7 (RL) | 4-5 (LR) | 5-6 (LR) | 6-7 (LR) | 4-6 (RLR) | 5-7 (RLR) | 4-6 (LRL) | 5-7 (LRL) | 4-7 (LRLR) |
|---|---|---|---|---|---|---|---|---|---|---|---|
| **Unliganded** | | | | | | | | | | | |
| **GPSGSGKS** | 0.24 | 1.99 | 0.26 | 1.42 | 0.11 | 2.24 | 0.00 | 0.36 | 0.29 | 0.01 | 0.03 |
| **GKPNVGKS** | 0.05 | 1.24 | 0.25 | 1.73 | 0.00 | 0.93 | 0.00 | 0.11 | 0.33 | 0.00 | 0.07 |
| **GEPGTGKS** | 0.05 | 1.10 | 0.23 | 1.65 | 0.02 | 2.46 | 0.00 | 0.21 | 0.21 | 0.00 | 0.03 |
| **GPTGSGKS** | 0.34 | 1.16 | 0.24 | 1.29 | 0.12 | 2.36 | 0.00 | 0.25 | 0.15 | 0.00 | 0.03 |
| **GPESSGKT** | 0.45 | 1.01 | 0.51 | 0.88 | 0.14 | 1.29 | 0.02 | 0.21 | 0.06 | 0.03 | 0.01 |
| **GVNGVGKT** | 0.01 | 1.04 | 0.24 | 1.36 | 0.01 | 2.22 | 0.00 | 0.14 | 0.12 | 0.00 | 0.00 |

**Table S11**. Occurrence (% simulation time) of correlated stretches of $\alpha_L$ and $\alpha_R$ conformations in Walker A-derived octapeptides in simulations with $C_{\alpha1}$-$C_{\alpha8}$ - restrained distances.

| **Sequence** | **4-5 (RL)** | **5-6 (RL)** | **6-7 (RL)** | **4-5 (LR)** | **5-6 (LR)** | **6-7 (LR)** | **4-6 (RLR)** | **5-7 (RLR)** | **4-6 (LRL)** | **5-7 (LRL)** | **4-7 (LRLR)** |
|---|---|---|---|---|---|---|---|---|---|---|---|
| **Unliganded** | | | | | | | | | | | |
| **GEPGTGKS** | 0.43 | 9.88 | 0.97 | 12.17 | 0.09 | 8.79 | 0.00 | 2.50 | 2.52 | 0.00 | 0.65 |
| **GPESSGKT** | 7.40 | 23.37 | 0.46 | 1.94 | 0.59 | 23.24 | 0.10 | 9.68 | 0.74 | 0.00 | 0.34 |
| **GVNGVGKT** | 0.05 | 11.16 | 0.78 | 12.49 | 0.03 | 11.63 | 0.00 | 2.30 | 4.24 | 0.00 | 1.20 |
| **$HPO_4^{2-}$** | | | | | | | | | | | |
| **GEPGTGKS** | 0.14 | 9.56 | 1.23 | 13.48 | 0.09 | 9.10 | 0.01 | 2.84 | 2.42 | 0.03 | 0.76 |
| **GPESSGKT** | 6.85 | 20.39 | 0.35 | 1.68 | 0.68 | 23.07 | 0.24 | 9.00 | 0.85 | 0.04 | 0.41 |
| **GVNGVGKT** | 0.66 | 8.68 | 0.90 | 16.13 | 0.08 | 13.66 | 0.00 | 2.64 | 4.22 | 0.01 | 1.81 |
| **GTP** | | | | | | | | | | | |
| **GEPGTGKS** | 0.00 | 2.53 | 0.01 | 70.66 | 0.00 | 0.83 | 0.00 | 0.56 | 1.75 | 0.00 | 0.46 |
| **GPESSGKT** | 8.55 | 10.31 | 0.39 | 4.06 | 0.82 | 25.85 | 0.57 | 4.22 | 1.25 | 0.00 | 0.62 |
| **GVNGVGKT** | 0.39 | 2.10 | 0.19 | 18.79 | 0.08 | 4.74 | 0.06 | 0.97 | 1.15 | 0.00 | 0.58 |

**Table S12**. Peptide-ligand hydrogen bonding (% simulation time) between the Walker A P-loop sequences studied in this work with $C_{\alpha 1}$-$C_{\alpha 8}$ distance restraint, in complex with either $HPO_4^{2-}$ or GTP.[a]

| | | $HPO_4^{2-}$ | | | GTP-α | | | GTP-β | | |
|---|---|---|---|---|---|---|---|---|---|---|
| **Sequence** | | GEPGTGKS | GPESSGKT | GVNGVGKT | GEPGTGKS | GPESSGKT | GVNGVGKT | GEPGTGKS | GPESSGKT | GVNGVGKT |
| | **N** | 11.11 | 10.78 | 13.78 | 41.11 | 40.48 | 45.06 | 39.77 | 36.20 | 29.17 |
| **1** | **R** | 0.00 | 0.00 | 0.00 | 0.00 | 0.00 | 0.00 | 0.00 | 0.00 | 0.00 |
| | **O** | 0.09 | 0.07 | 0.09 | 0.00 | 0.00 | 0.00 | 0.00 | 0.00 | 0.00 |
| | **N** | 1.11 | 0.00 | 3.18 | 1.12 | 0.00 | 13.47 | 0.03 | 0.00 | 0.48 |
| **2** | **R** | 0.01 | 0.00 | 0.00 | 0.00 | 0.00 | 0.00 | 0.00 | 0.00 | 0.00 |
| | **O** | 0.10 | 0.01 | 0.06 | 0.00 | 0.00 | 0.00 | 0.00 | 0.00 | 0.00 |
| | **N** | 0.00 | 0.20 | 3.36 | 0.00 | 0.00 | 15.26 | 0.00 | 0.00 | 0.90 |
| **3** | **R** | 0.00 | 0.02 | 3.90 | 0.00 | 0.00 | 13.12 | 0.00 | 0.00 | 0.65 |
| | **O** | 0.00 | 0.00 | 0.07 | 0.00 | 0.00 | 0.00 | 0.00 | 0.00 | 0.00 |
| | **N** | 0.19 | 0.21 | 1.72 | 0.01 | 0.00 | 10.29 | 0.00 | 0.00 | 0.31 |
| **4** | **R** | 0.00 | 4.86 | 0.00 | 0.00 | 15.62 | 0.00 | 0.00 | 2.04 | 0.00 |
| | **O** | 0.01 | 0.01 | 0.01 | 0.00 | 0.00 | 0.00 | 0.00 | 0.00 | 0.00 |
| | **N** | 0.48 | 0.33 | 0.66 | 0.01 | 0.03 | 6.30 | 0.00 | 0.00 | 0.15 |
| **5** | **R** | 0.75 | 4.54 | 0.00 | 0.58 | 4.03 | 0.00 | 0.03 | 2.66 | 0.00 |
| | **O** | 0.01 | 0.01 | 0.00 | 0.00 | 0.00 | 0.00 | 0.00 | 0.00 | 0.00 |
| | **N** | 0.32 | 0.30 | 0.61 | 0.01 | 2.62 | 6.75 | 0.00 | 0.19 | 0.16 |
| **6** | **R** | 0.00 | 0.00 | 0.00 | 0.00 | 0.00 | 0.00 | 0.00 | 0.00 | 0.00 |
| | **O** | 0.02 | 0.01 | 0.02 | 0.00 | 0.00 | 0.00 | 0.00 | 0.00 | 0.00 |
| | **N** | 0.30 | 0.25 | 0.85 | 0.47 | 7.65 | 15.87 | 0.06 | 0.79 | 7.46 |
| **7** | **R** | 15.67 | 14.42 | 15.10 | 37.75 | 37.81 | 47.66 | 29.91 | 30.11 | 34.65 |
| | **O** | 0.01 | 0.00 | 0.01 | 0.00 | 0.00 | 0.00 | 0.00 | 0.00 | 0.00 |
| | **N** | 0.47 | 0.29 | 0.82 | 0.03 | 3.12 | 5.54 | 0.04 | 0.15 | 0.10 |
| **8** | **R** | 1.37 | 0.41 | 1.06 | 0.05 | 3.38 | 6.70 | 0.06 | 1.08 | 0.92 |
| | **O** | 0.01 | 0.00 | 0.01 | 0.00 | 0.00 | 0.00 | 0.00 | 0.00 | 0.00 |

| Sequence | | GTP-γ | | | GTP-triphosphate | | | GTP base+sugar | | |
|---|---|---|---|---|---|---|---|---|---|---|
| | | GEPGTGKS | GPESSGKT | GVNGVGKT | GEPGTGKS | GPESSGKT | GVNGVGKT | GEPGTGKS | GPESSGKT | GVNGVGKT |
| | **N** | 51.80 | 36.19 | 53.10 | 81.67 | 76.05 | 81.63 | 80.15 | 43.27 | 47.25 |
| **1** | **R** | 0.00 | 0.00 | 0.00 | 0.00 | 0.00 | 0.00 | 0.00 | 0.00 | 0.00 |
| | **O** | 0.00 | 0.00 | 0.00 | 0.00 | 0.00 | 0.00 | 0.17 | 0.32 | 1.99 |
| | **N** | 0.16 | 0.00 | 7.98 | 1.31 | 0.00 | 21.83 | 0.45 | 0.00 | 1.42 |
| **2** | **R** | 0.00 | 0.00 | 0.00 | 0.00 | 0.00 | 0.00 | 0.17 | 0.00 | 0.00 |
| | **O** | 0.00 | 0.00 | 0.00 | 0.00 | 0.00 | 0.00 | 1.65 | 0.36 | 2.10 |
| | **N** | 0.00 | 0.04 | 8.05 | 0.00 | 0.05 | 23.96 | 0.00 | 1.19 | 1.68 |
| **3** | **R** | 0.00 | 0.00 | 8.25 | 0.00 | 0.00 | 21.35 | 0.00 | 1.85 | 4.67 |
| | **O** | 0.00 | 0.00 | 0.00 | 0.00 | 0.00 | 0.00 | 0.01 | 0.27 | 15.92 |
| | **N** | 0.00 | 0.08 | 2.17 | 0.01 | 0.09 | 12.50 | 15.69 | 2.11 | 12.62 |
| **4** | **R** | 0.00 | 12.82 | 0.00 | 0.00 | 30.31 | 0.00 | 0.00 | 2.06 | 0.00 |
| | **O** | 0.00 | 0.00 | 0.00 | 0.00 | 0.00 | 0.00 | 0.06 | 1.15 | 17.69 |
| | **N** | 0.00 | 0.01 | 0.00 | 0.01 | 0.04 | 6.45 | 32.33 | 0.88 | 5.61 |
| **5** | **R** | 0.07 | 5.40 | 0.00 | 0.68 | 11.89 | 0.00 | 6.87 | 2.97 | 0.00 |
| | **O** | 0.00 | 0.00 | 0.00 | 0.00 | 0.00 | 0.00 | 19.72 | 2.51 | 2.32 |
| | **N** | 0.00 | 0.01 | 0.00 | 0.01 | 2.77 | 6.82 | 47.38 | 2.81 | 10.76 |
| **6** | **R** | 0.00 | 0.00 | 0.00 | 0.00 | 0.00 | 0.00 | 0.00 | 0.00 | 0.00 |
| | **O** | 0.00 | 0.00 | 0.00 | 0.00 | 0.00 | 0.00 | 68.99 | 3.24 | 8.55 |
| | **N** | 0.05 | 0.45 | 1.07 | 0.58 | 8.76 | 20.25 | 2.39 | 1.67 | 10.28 |
| **7** | **R** | 82.19 | 49.26 | 64.87 | 91.94 | 67.37 | 82.01 | 46.77 | 5.94 | 9.18 |
| | **O** | 0.00 | 0.00 | 0.00 | 0.00 | 0.00 | 0.00 | 0.47 | 2.11 | 3.73 |
| | **N** | 1.24 | 0.42 | 1.05 | 1.30 | 3.67 | 6.68 | 0.39 | 2.90 | 7.23 |
| **8** | **R** | 1.54 | 0.54 | 1.26 | 1.65 | 4.75 | 8.81 | 1.26 | 4.25 | 7.92 |
| | **O** | 0.00 | 0.00 | 0.00 | 0.00 | 0.00 | 0.00 | 2.91 | 15.27 | 24.62 |

[a]Analysis based on snapshots taken every 10 ps of the simulation, calculated as described in the main text. This data is shown visually in **Figure S7**.

**Table S13**. Occurrence (% simulation time) of correlated stretches of $\alpha_L$ and $\alpha_R$ conformations in control octapeptides.

| **Sequence** | **4-5 (RL)** | **5-6 (RL)** | **6-7 (RL)** | **4-5 (LR)** | **5-6 (LR)** | **6-7 (LR)** | **4-6 (RLR)** | **5-7 (RLR)** | **4-6 (LRL)** | **5-7 (LRL)** | **4-7 (LRLR)** |
|---|---|---|---|---|---|---|---|---|---|---|---|
| **Unliganded** | | | | | | | | | | | |
| **KSPGGSSG** | 1.23 | 0.94 | 3.75 | 1.07 | 5.05 | 3.83 | 0.51 | 0.35 | 0.26 | 0.18 | 0.05 |
| **KGSGNKVP** | 0.73 | 2.01 | 0.25 | 3.44 | 4.85 | 0.07 | 0.56 | 0.00 | 0.57 | 0.01 | 0.00 |
| **EGKPGTSG** | 0.40 | 0.17 | 6.50 | 0.00 | 19.13 | 1.23 | 0.21 | 0.04 | 0.00 | 2.74 | 0.00 |
| **PGKGGSST** | 0.49 | 0.64 | 8.83 | 0.54 | 4.70 | 3.94 | 0.12 | 0.24 | 0.05 | 0.70 | 0.02 |
| **TGPSSEKG** | 3.44 | 0.73 | 1.16 | 4.58 | 8.37 | 2.13 | 0.97 | 0.13 | 0.04 | 0.00 | 0.01 |
| **VVTGGNKG** | 0.76 | 0.70 | 1.08 | 1.38 | 3.91 | 2.31 | 0.11 | 0.13 | 0.09 | 0.23 | 0.01 |
| **$HPO_4^{2-}$** | | | | | | | | | | | |
| **KSPGGSSG** | 1.03 | 0.81 | 2.34 | 1.21 | 5.79 | 4.27 | 0.24 | 0.37 | 0.03 | 0.30 | 0.01 |
| **KGSGNKVP** | 0.80 | 1.97 | 0.19 | 4.06 | 6.52 | 0.01 | 0.23 | 0.00 | 0.29 | 0.08 | 0.00 |
| **EGKPGTSG** | 0.39 | 0.13 | 6.11 | 0.00 | 17.87 | 0.79 | 0.18 | 0.03 | 0.00 | 2.52 | 0.00 |
| **PGKGGSST** | 0.02 | 1.22 | 1.50 | 0.09 | 0.53 | 9.65 | 0.00 | 0.49 | 0.03 | 0.07 | 0.02 |
| **TGPSSEKG** | 4.66 | 0.77 | 1.51 | 5.70 | 5.68 | 3.54 | 1.30 | 0.16 | 0.29 | 0.01 | 0.03 |
| **VVTGGNKG** | 0.78 | 0.34 | 0.91 | 1.56 | 3.86 | 1.08 | 0.12 | 0.12 | 0.03 | 0.07 | 0.01 |
| **GTP** | | | | | | | | | | | |
| **KSPGGSSG** | 0.00 | 0.58 | 9.96 | 0.00 | 1.11 | 0.50 | 0.00 | 0.06 | 0.00 | 0.54 | 0.00 |
| **KGSGNKVP** | 7.04 | 0.56 | 0.00 | 0.02 | 4.33 | 0.07 | 1.18 | 0.00 | 0.00 | 0.00 | 0.00 |
| **EGKPGTSG** | 0.00 | 23.50 | 0.02 | 1.16 | 0.15 | 6.81 | 0.00 | 4.19 | 0.27 | 0.00 | 0.00 |
| **PGKGGSST** | 0.08 | 2.81 | 1.16 | 0.33 | 0.45 | 12.06 | 0.00 | 0.69 | 0.10 | 0.01 | 0.04 |
| **TGPSSEKG** | 5.54 | 18.19 | 0.37 | 2.96 | 0.29 | 10.67 | 0.29 | 2.86 | 1.18 | 0.24 | 0.07 |
| **VVTGGNKG** | 3.68 | 0.99 | 5.56 | 0.63 | 3.73 | 0.13 | 2.18 | 0.00 | 0.08 | 0.94 | 0.00 |

**Table S14**. Peptide-ligand hydrogen bonding (% simulation time) between the control sequences studied in this work, in complex with either $HPO_4^{2-}$ or GTP.[a]

| Sequence | | $HPO_4^{2-}$ KSPGGSSG | $HPO_4^{2-}$ KGSGNKVP | $HPO_4^{2-}$ EGKPGTSG | $HPO_4^{2-}$ PGKGGSST | $HPO_4^{2-}$ TGPSSEKG | $HPO_4^{2-}$ VVTGGNKG | GTP-$\alpha$ KSPGGSSG | GTP-$\alpha$ KGSGNKVP | GTP-$\alpha$ EGKPGTSG | GTP-$\alpha$ PGKGGSST | GTP-$\alpha$ TGPSSEKG | GTP-$\alpha$ VVTGGNKG |
|---|---|---|---|---|---|---|---|---|---|---|---|---|---|
| | **N** | 24.72 | 23.02 | 11.33 | 19.93 | 13.01 | 10.87 | 51.75 | 20.66 | 46.54 | 43.96 | 52.64 | 42.45 |
| **1** | **R** | 14.93 | 15.36 | 0.02 | 0.00 | 4.44 | 0.00 | 19.64 | 25.40 | 0.00 | 0.00 | 12.57 | 0.00 |
| | **O** | 0.11 | 0.11 | 0.12 | 0.21 | 0.10 | 0.06 | 0.00 | 0.00 | 0.00 | 0.00 | 0.00 | 0.00 |
| | **N** | 16.49 | 13.62 | 3.08 | 8.09 | 5.95 | 2.16 | 27.25 | 0.06 | 10.08 | 22.79 | 7.65 | 0.21 |
| **2** | **R** | 23.06 | 0.00 | 0.00 | 0.00 | 0.00 | 0.00 | 27.22 | 0.00 | 0.00 | 0.00 | 0.00 | 0.00 |
| | **O** | 0.05 | 0.12 | 0.09 | 0.22 | 0.14 | 0.01 | 0.00 | 0.00 | 0.00 | 0.00 | 0.00 | 0.00 |
| | **N** | 0.00 | 11.10 | 0.40 | 1.73 | 0.00 | 3.65 | 0.00 | 39.27 | 3.80 | 12.82 | 0.00 | 3.77 |
| **3** | **R** | 0.00 | 11.76 | 12.41 | 15.88 | 0.00 | 3.60 | 0.00 | 35.69 | 34.86 | 40.75 | 0.00 | 24.59 |
| | **O** | 0.00 | 0.02 | 0.02 | 0.05 | 0.00 | 0.03 | 0.00 | 0.00 | 0.00 | 0.00 | 0.00 | 0.00 |
| | **N** | 0.14 | 3.20 | 0.00 | 0.73 | 0.47 | 1.83 | 0.01 | 8.42 | 0.00 | 5.09 | 0.17 | 3.14 |
| **4** | **R** | 0.00 | 0.00 | 0.00 | 0.00 | 2.73 | 0.00 | 0.00 | 0.00 | 0.00 | 0.00 | 2.55 | 0.00 |
| | **O** | 0.01 | 0.06 | 0.00 | 0.04 | 0.01 | 0.03 | 0.00 | 0.00 | 0.00 | 0.00 | 0.00 | 0.00 |
| | **N** | 0.09 | 0.40 | 0.09 | 0.24 | 0.39 | 0.45 | 0.87 | 2.80 | 0.03 | 1.44 | 0.61 | 3.95 |
| **5** | **R** | 0.00 | 1.83 | 0.00 | 0.00 | 3.84 | 0.00 | 0.00 | 3.49 | 0.00 | 0.00 | 8.65 | 0.00 |
| | **O** | 0.13 | 0.02 | 0.00 | 0.01 | 0.01 | 0.01 | 0.00 | 0.00 | 0.00 | 0.00 | 0.00 | 0.00 |
| | **N** | 2.60 | 0.51 | 0.56 | 1.85 | 0.05 | 0.35 | 1.87 | 3.27 | 0.05 | 2.73 | 0.26 | 1.92 |
| **6** | **R** | 6.13 | 14.50 | 0.90 | 4.28 | 0.01 | 0.99 | 2.16 | 24.78 | 0.98 | 2.90 | 0.00 | 6.08 |
| | **O** | 0.02 | 0.00 | 0.01 | 0.01 | 0.01 | 0.01 | 0.00 | 0.00 | 0.00 | 0.00 | 0.00 | 0.00 |
| | **N** | 2.90 | 0.08 | 0.62 | 1.68 | 0.03 | 0.13 | 1.26 | 0.01 | 0.04 | 2.14 | 1.87 | 2.07 |
| **7** | **R** | 5.68 | 0.00 | 2.49 | 3.34 | 14.76 | 14.47 | 2.73 | 0.00 | 1.06 | 3.37 | 29.03 | 49.52 |
| | **O** | 0.01 | 0.01 | 0.01 | 0.00 | 0.01 | 0.00 | 0.00 | 0.00 | 0.00 | 0.00 | 0.00 | 0.00 |
| | **N** | 0.43 | 0.00 | 0.14 | 0.66 | 0.07 | 0.03 | 0.49 | 0.00 | 0.24 | 0.09 | 1.84 | 0.20 |
| **8** | **R** | 0.00 | 0.00 | 0.00 | 0.89 | 0.00 | 0.00 | 0.00 | 0.00 | 0.00 | 0.31 | 0.00 | 0.00 |
| | **O** | 0.02 | 0.01 | 0.01 | 0.01 | 0.01 | 0.01 | 0.00 | 0.00 | 0.00 | 0.00 | 0.00 | 0.00 |

| Sequence | | GTP-β | | | | | | GTP-γ | | | | | |
|---|---|---|---|---|---|---|---|---|---|---|---|---|---|
| | | KSPGGSSG | KGSGNKVP | EGKPGTSG | PGKGGSST | TGPSSEKG | VVTGGNKG | KSPGGSSG | KGSGNKVP | EGKPGTSG | PGKGGSST | TGPSSEKG | VVTGGNKG |
| | **N** | 20.49 | 17.22 | 26.01 | 18.44 | 24.45 | 20.95 | 67.04 | 43.86 | 47.28 | 39.13 | 45.74 | 23.56 |
| **1** | **R** | 20.86 | 27.55 | 0.00 | 0.00 | 2.70 | 0.00 | 33.01 | 46.30 | 0.00 | 0.00 | 20.03 | 0.00 |
| | **O** | 0.00 | 0.00 | 0.00 | 0.00 | 0.00 | 0.00 | 0.00 | 0.00 | 0.00 | 0.00 | 0.00 | 0.00 |
| | **N** | 6.46 | 0.08 | 3.25 | 10.83 | 4.49 | 0.17 | 35.74 | 0.01 | 15.53 | 14.89 | 27.04 | 0.05 |
| **2** | **R** | 5.49 | 0.00 | 0.00 | 0.00 | 0.00 | 0.00 | 40.42 | 0.00 | 0.00 | 0.00 | 0.00 | 0.00 |
| | **O** | 0.00 | 0.00 | 0.00 | 0.00 | 0.00 | 0.00 | 0.00 | 0.00 | 0.00 | 0.00 | 0.00 | 0.00 |
| | **N** | 0.00 | 7.05 | 1.31 | 8.42 | 0.00 | 0.92 | 0.00 | 6.17 | 3.85 | 2.98 | 0.00 | 0.93 |
| **3** | **R** | 0.00 | 8.27 | 21.69 | 33.44 | 0.00 | 7.58 | 0.00 | 7.86 | 38.21 | 51.76 | 0.00 | 2.93 |
| | **O** | 0.00 | 0.00 | 0.00 | 0.00 | 0.00 | 0.00 | 0.00 | 0.00 | 0.00 | 0.00 | 0.00 | 0.00 |
| | **N** | 0.00 | 0.40 | 0.00 | 1.36 | 0.00 | 0.27 | 0.00 | 0.47 | 0.00 | 0.48 | 1.52 | 0.60 |
| **4** | **R** | 0.00 | 0.00 | 0.00 | 0.00 | 0.82 | 0.00 | 0.00 | 0.00 | 0.00 | 0.00 | 5.21 | 0.00 |
| | **O** | 0.00 | 0.00 | 0.00 | 0.00 | 0.00 | 0.00 | 0.00 | 0.00 | 0.00 | 0.00 | 0.00 | 0.00 |
| | **N** | 0.06 | 0.26 | 0.00 | 0.10 | 0.05 | 0.66 | 0.01 | 0.05 | 0.01 | 0.22 | 1.21 | 0.51 |
| **5** | **R** | 0.00 | 0.69 | 0.00 | 0.00 | 2.64 | 0.00 | 0.00 | 1.04 | 0.00 | 0.00 | 3.34 | 0.00 |
| | **O** | 0.00 | 0.00 | 0.00 | 0.00 | 0.00 | 0.00 | 0.00 | 0.00 | 0.00 | 0.00 | 0.00 | 0.00 |
| | **N** | 0.07 | 0.07 | 0.00 | 0.46 | 0.02 | 0.03 | 0.08 | 0.22 | 0.00 | 0.73 | 0.04 | 0.04 |
| **6** | **R** | 0.13 | 19.21 | 5.04 | 0.86 | 0.00 | 1.39 | 0.97 | 31.80 | 1.93 | 2.54 | 0.00 | 0.73 |
| | **O** | 0.00 | 0.00 | 0.00 | 0.00 | 0.00 | 0.00 | 0.00 | 0.00 | 0.00 | 0.00 | 0.00 | 0.00 |
| | **N** | 0.01 | 0.00 | 0.00 | 0.42 | 0.00 | 0.54 | 0.08 | 0.00 | 0.16 | 1.82 | 0.00 | 0.00 |
| **7** | **R** | 0.43 | 0.00 | 0.20 | 1.01 | 25.73 | 26.82 | 1.67 | 0.00 | 0.64 | 3.06 | 58.01 | 58.74 |
| | **O** | 0.00 | 0.00 | 0.00 | 0.00 | 0.00 | 0.00 | 0.00 | 0.00 | 0.00 | 0.00 | 0.00 | 0.00 |
| | **N** | 0.02 | 0.00 | 0.17 | 0.01 | 0.37 | 0.00 | 0.01 | 0.00 | 0.03 | 0.00 | 0.00 | 0.00 |
| **8** | **R** | 0.00 | 0.00 | 0.00 | 0.10 | 0.00 | 0.00 | 0.00 | 0.00 | 0.00 | 0.06 | 0.00 | 0.00 |
| | **O** | 0.00 | 0.00 | 0.00 | 0.00 | 0.00 | 0.00 | 0.00 | 0.00 | 0.00 | 0.00 | 0.00 | 0.00 |

| Sequence | | GTP-triphosphate | | | | | | GTP base+sugar | | | | | |
|---|---|---|---|---|---|---|---|---|---|---|---|---|---|
| | | KSPGGSSG | KGSGNKVP | EGKPGTSG | PGKGGSST | TGPSSEKG | VVTGGNKG | KSPGGSSG | KGSGNKVP | EGKPGTSG | PGKGGSST | TGPSSEKG | VVTGGNKG |
| | **N** | 86.08 | 63.24 | 74.53 | 73.00 | 79.28 | 62.33 | 41.68 | 15.59 | 21.29 | 7.79 | 63.99 | 29.68 |
| **1** | **R** | 50.55 | 62.57 | 0.00 | 0.00 | 34.98 | 0.00 | 2.01 | 3.24 | 3.10 | 0.00 | 0.19 | 0.00 |
| | **O** | 0.00 | 0.00 | 0.00 | 0.00 | 0.00 | 0.00 | 1.29 | 0.50 | 3.25 | 3.39 | 1.06 | 1.20 |
| | **N** | 67.65 | 0.14 | 27.88 | 42.52 | 38.74 | 0.42 | 6.93 | 3.92 | 8.64 | 6.40 | 1.58 | 2.22 |
| **2** | **R** | 72.37 | 0.00 | 0.00 | 0.00 | 0.00 | 0.00 | 3.81 | 0.00 | 0.00 | 0.00 | 0.00 | 0.00 |
| | **O** | 0.00 | 0.00 | 0.00 | 0.00 | 0.00 | 0.00 | 4.75 | 8.17 | 4.60 | 5.36 | 0.47 | 4.17 |
| | **N** | 0.00 | 51.15 | 8.90 | 23.29 | 0.00 | 5.57 | 0.00 | 3.69 | 5.03 | 4.69 | 0.00 | 0.82 |
| **3** | **R** | 0.00 | 51.68 | 58.74 | 72.46 | 0.00 | 34.66 | 0.00 | 3.90 | 20.19 | 9.75 | 0.00 | 3.44 |
| | **O** | 0.00 | 0.00 | 0.00 | 0.00 | 0.00 | 0.00 | 2.31 | 1.92 | 5.33 | 3.00 | 2.03 | 2.77 |
| | **N** | 0.01 | 9.18 | 0.00 | 5.89 | 1.69 | 3.97 | 1.38 | 13.52 | 0.00 | 3.48 | 6.47 | 6.29 |
| **4** | **R** | 0.00 | 0.00 | 0.00 | 0.00 | 8.50 | 0.00 | 0.00 | 0.00 | 0.00 | 0.00 | 0.63 | 0.00 |
| | **O** | 0.00 | 0.00 | 0.00 | 0.00 | 0.00 | 0.00 | 8.07 | 14.51 | 7.42 | 4.18 | 16.68 | 7.36 |
| | **N** | 0.94 | 2.94 | 0.03 | 1.71 | 1.86 | 4.69 | 4.60 | 7.42 | 1.32 | 6.77 | 21.86 | 4.16 |
| **5** | **R** | 0.00 | 4.95 | 0.00 | 0.00 | 14.44 | 0.00 | 0.00 | 4.56 | 0.00 | 0.00 | 8.14 | 0.00 |
| | **O** | 0.00 | 0.00 | 0.00 | 0.00 | 0.00 | 0.00 | 5.68 | 11.40 | 6.50 | 6.82 | 0.77 | 12.34 |
| | **N** | 1.98 | 3.54 | 0.05 | 3.79 | 0.32 | 1.99 | 5.10 | 7.29 | 1.03 | 8.59 | 38.85 | 4.12 |
| **6** | **R** | 3.23 | 50.28 | 7.71 | 6.21 | 0.00 | 7.75 | 2.90 | 6.78 | 4.10 | 3.84 | 20.64 | 8.12 |
| | **O** | 0.00 | 0.00 | 0.00 | 0.00 | 0.00 | 0.00 | 1.67 | 4.03 | 1.48 | 2.46 | 1.84 | 4.66 |
| | **N** | 1.34 | 0.01 | 0.21 | 4.14 | 1.87 | 2.61 | 6.22 | 1.48 | 1.30 | 8.53 | 1.54 | 4.83 |
| **7** | **R** | 4.80 | 0.00 | 1.88 | 7.28 | 71.57 | 76.88 | 4.31 | 0.00 | 2.94 | 4.16 | 3.62 | 11.06 |
| | **O** | 0.00 | 0.00 | 0.00 | 0.00 | 0.00 | 0.00 | 1.84 | 2.56 | 3.18 | 4.43 | 1.25 | 2.38 |
| | **N** | 0.51 | 0.00 | 0.39 | 0.09 | 2.17 | 0.21 | 5.24 | 0.00 | 5.45 | 2.35 | 1.62 | 2.81 |
| **8** | **R** | 0.00 | 0.00 | 0.00 | 0.47 | 0.00 | 0.00 | 0.00 | 0.00 | 0.00 | 5.57 | 0.00 | 0.00 |
| | **O** | 0.00 | 0.00 | 0.00 | 0.00 | 0.00 | 0.00 | 17.33 | 33.95 | 22.91 | 14.70 | 44.40 | 19.44 |

[a]Analysis based on snapshots taken every 10 ps of the simulation, calculated as described in the main text. This data is shown visually in **Figure S9**.

**Table S15.** List of distance restraints between the $C_{\alpha 1}$ and $C_{\alpha 8}$ atoms of the peptides studied in the restrained simulations.[a]

| **Sequence** | **$r_0$, nm** | **$r_1$, nm** | **$r_2$, nm** | **Force constant, kJ mol$^{-1}$ nm$^{-1}$** |
|---|---|---|---|---|
| GEPGTGKS | 1.00 | 1.11 | 1.20 | 4184 |
| GPESSGKT | 0.94 | 01.04 | 1.14 | 4184 |
| GVNGVGKT | 0.94 | 01.04 | 1.14 | 4184 |

[a] A piecewise linear-harmonict restrain potential was used (bond type 10 in GROMACS (15)). The potential is quadratic for $r_{ij} \leq r_0$, $r_0 \leq r_{ij} \leq r_1$ and $r_1 \leq r_{ij} \leq r_2$, and linear for $r_2 \leq r_{ij}$.
